# Single-cell ATAC and RNA sequencing reveal pre-existing and persistent subpopulations of cells associated with relapse of prostate cancer

**DOI:** 10.1101/2021.02.09.430114

**Authors:** S Taavitsainen, N Engedal, S Cao, F Handle, A Erickson, S Prekovic, D Wetterskog, T Tolonen, EM Vuorinen, A Kiviaho, R Nätkin, T Häkkinen, W Devlies, S Henttinen, R Kaarijärvi, M Lahnalampi, H Kaljunen, K Nowakowska, H Syvälä, M Bläuer, P Cremaschi, F Claessens, T Visakorpi, TLJ Tammela, T Murtola, KJ Granberg, AD Lamb, K Ketola, IG Mills, G Attard, W Wang, M Nykter, A Urbanucci

## Abstract

Prostate cancer is profoundly heterogeneous and patients would benefit from methods that stratify clinically indolent from more aggressive forms of the disease. We employed single-cell assay for transposase-accessible chromatin (ATAC) and RNA sequencing in models of early treatment response and resistance to enzalutamide. In doing so, we identified pre-existing and treatment-persistent cell subpopulations that possess transcriptional stem-like features and regenerative potential when subjected to treatment. We found distinct chromatin landscapes associated with enzalutamide treatment and resistance that are linked to alternative transcriptional programs. Transcriptional profiles characteristic of persistent stem-like cells were able to stratify the treatment response of patients. Ultimately, we show that defining changes in chromatin and gene expression in single-cell populations from pre-clinical models can reveal hitherto unrecognized molecular predictors of treatment response. This suggests that high analytical resolution of pre-clinical models may powerfully inform clinical decision-making.

## Introduction

Prostate cancer (PC) relies on androgen receptor (AR) signaling for development and progression. Progression on androgen deprivation therapy (ADT) or AR signaling inhibitors (ARSIs) leads to castration resistant (CRPC) or treatment-induced neuroendocrine prostate cancer (NEPC) (Beltran et al., 2016). The most frequently characterized mechanisms of PC or CRPC resistance to ARSIs, ADT, or both, revolve around re-establishing AR signaling e.g. via overexpression of AR or *AR* mutations (Abida et al., 2019; Alumkal et al., 2020; Devlies et al., 2020; He et al., 2018). Additionally, forms of resistance that are indifferent to AR (Handle et al., 2019), more dependent on FGF signaling (Bluemn et al., 2017) or that are AR negative with NEPC-like features (Beltran et al., 2014) are being identified.

PC is profoundly heterogeneous (Haffner et al., 2020; Løvf et al., 2019; Tomlins et al., 2015; Woodcock et al., 2020) and patients would benefit from methods that differentiate between clinically mild disease and more aggressive forms. To date, most studies characterizing genetic mutations causing drug resistance (Gerhauser et al., 2018; Grasso et al., 2012; Taylor et al., 2010) point at the selection of a single dominant clone to identify molecular predictors (Haffner et al., 2020). However, non-genetic effects linked to transcriptomic changes, although more common (Abida et al., 2019; Alumkal et al., 2020; Devlies et al., 2020; He et al., 2018), are less understood. Moreover, most RNA sequencing data has been obtained from the bulk of the tumors, which cannot account for PC heterogeneity. The tumor transcriptome is the result of several biological processes contributing to differential gene regulation that is not necessarily in sync in all cells within the bulk (Su et al., 2020; Zhang et al., 2020). Non-genetic effects linked to epigenetics in PC drug resistance are even less understood. Chromatin structure and DNA accessibility to transcription factor (TF) DNA binding motifs is the first layer of gene regulation (Sönmezer et al., 2020; Strickfaden et al., 2020). For example, expression and activity of the Glucocorticoid Receptor and the pluripotent stem cell transcription factor Sox2 have been shown to promote enzalutamide (ENZ) resistance (Crona and Whang, 2017; Isikbay et al., 2014; Li et al., 2017a; Mu et al., 2017). Moreover, exposure to ENZ can alter activity of other chromatin regulators (Beltran et al., 2016).

Only a few studies to date have used single-cell sequencing technology on human PC tumor specimens (Chen et al., 2021; Cyrta et al., 2020; Dong et al., 2020; He et al.; Karthaus et al., 2020; Ma et al., 2020b; Song et al.). Although these approaches characterize features of individual tumors, they do not allow for elucidation of the temporal sequence of events occurring during the evolution of drug resistance. To explore how heterogeneous PC cells respond to ARSIs, we analyzed the evolution of resistance in the epithelial-derived component of PC in models of ENZ exposed and resistant PC cell lines at a single-cell level. Through deconvolution of transcriptional signals from molecular gene classifiers derived in this study, we show evidence of treatment-persistent and pre-existing PC cells that can predict treatment response in both primary and advanced patients.

## Results

### Chromatin reprogramming underpins enzalutamide resistance

To study molecular consequences of AR signaling suppression and drug resistance dynamics in PC, we used LNCaP parental and LNCaP-derived ENZ-resistant cell lines RES-A and RES-B generated via long-term exposure to AR-targeting agents (Handle et al., 2019)(see **Methods**), as well as other independently generated LNCaP and VCaP-derived models (**Figure 1A**). Both castration- and ENZ-resistant prostate cancer are characterized by increased AR signaling (Formaggio et al., 2021; Watson et al., 2015). Along with others, we previously reported that AR overexpression is associated with chromatin reprogramming (Braadland et al., 2019; Jia et al., 2006; Tewari et al., 2012; Urbanucci et al., 2017). As transcriptionally permissive chromatin in open conformation is the first layer of gene regulation (Sönmezer et al., 2020; Strickfaden et al., 2020), we hypothesized that chromatin structure would be reshaped in ENZ-resistant cells and lead to modification of the transcriptome.

**Figure 1.**
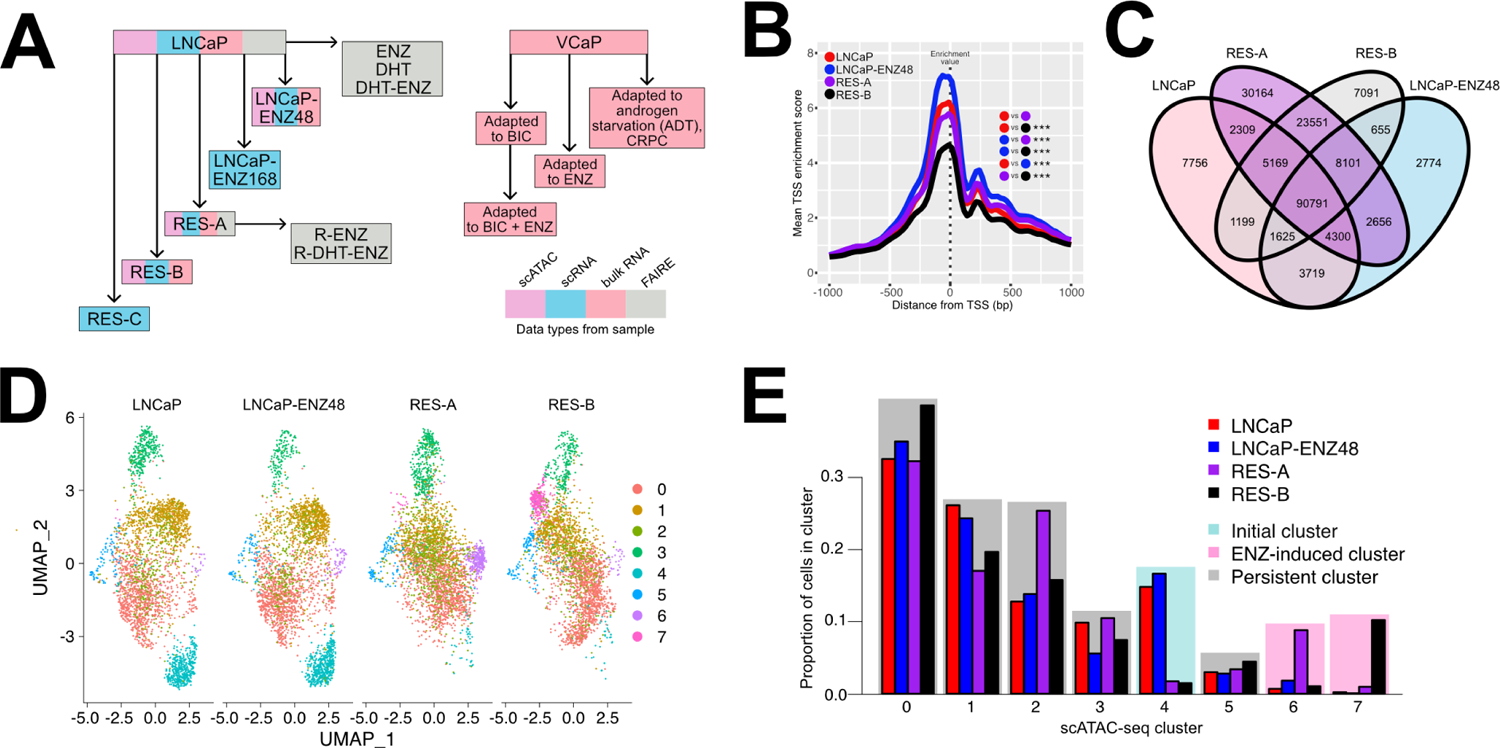
Chromatin reprogramming in enzalutamide resistance. **(A)** Overview of the cell lines models, assays, and treatments included in the study. Boxes with sample names are colored according to the data types generated from the sample (single-cell ATAC-, single-cell RNA-, bulk RNA-, and/or FAIRE-sequencing). (**B**) LNCaP parental, LNCaP-ENZ48, RES-A, and RES-B single-cell (sc) ATAC-seq enrichment score in a 2kb window around the transcription start site (TSS). Enrichment scores at each TSS (position 0 in the plot) were used as the enrichment values and compared between pairs of samples.

To extrapolate the contribution of chromatin structure to ENZ resistance, we performed single-cell (sc) assays for transposase-accessible chromatin and sequencing (scATAC-seq) on four samples: (1) LNCaP parental cells (LNCaP), (2) LNCaP exposed to short-term (48 hours) ENZ (10 μM) treatment (LNCaP-ENZ48), (3) RES-A and (4) RES-B (**Figure 1A**). We first analyzed the scATAC-seq data as if it would have been sequenced in bulk cells (see **Methods**). The ATAC-seq signal at transcription start sites (TSS) decreased in ENZ-resistant cells compared to parental, particularly in RES-B cells (average enrichment score 4.8 in RES-B vs 6.2 in LNCaP, p < 0.001, *t*-test) (**Figure 1B**). RES-A and RES-B cells shared a large proportion (14% in RES-A and 17% in RES-B) of “ENZ-resistant-specific” open chromatin regions not found in parental LNCaP. Additionally, RES-A cells had a higher proportion of unique open sites compared to RES-B (19% vs 5%, *P* < 0.001, chi-square test) and LNCaP (19% vs 7%, *P* < 0.001, chi-square test) (**Figure 1C**). These findings are consistent with TSS non-targeted opening (Jiang and Zhang, 2021) of the chromatin upon enzalutamide resistance.

We confirmed the extent of chromatin opening and reprogramming in ENZ-resistant cells by performing formaldehyde-assisted isolation of regulatory elements (FAIRE) sequencing (Giresi et al., 2007) on the parental LNCaP and RES-A cells subjected to androgen starvation, or exposed to androgens, ENZ, or both agents (**Figure S1A-D**)(see **Methods**). Even in this bulk assay, ENZ and androgen starvation appeared to be more significant drivers of reprogramming in RES-A than in parental LNCaP. While there was no difference in the total number of open chromatin sites, ENZ-resistant samples had a higher proportion of unique open sites compared to parental in the presence of androgens (24% vs 12%, *P* < 0.001, chi-square test) (**Figure S1E**) and in androgen-deprived (castrate) conditions (27% vs 9%, *P* < 0.001, chi-square test) (**Figure S1C**). Read distribution analysis (see **Methods)** demonstrated that the chromatin of ENZ-resistant cells is more open in castrate conditions (p = 0.018, *t*-test) (**Figure S1D**) but not in presence of androgens (p = 0.239, *t*-test) (**Figure S1F**), and that ENZ has an additive effect on castration in inducing chromatin compaction in parental ENZ-sensitive cells that is counteracted by androgens (**Figure S1D, Figure S1F**).

Next, we used all samples with scATAC-seq to generate cluster visualizations of cell subpopulations with different chromatin (see **Methods**) (**Figure 1D**). We identified clusters which we termed “unique” or “shared” across the samples (**Figure 1E**). Unique clusters were specific to RES-A, RES-B, or both (named “ENZ-induced clusters”), or specific to the untreated and/or short-term ENZ-treated parental line (named “initial clusters”). Shared clusters were present at similar proportions across the samples and were named “persistent clusters” (**Figure 1E**). We compared each cluster to all other clusters to determine its unique chromatin profile based on differentially accessible chromatin regions (DARs).

The most prevalent chromatin-based scATAC-seq clusters (0, 1, and 2) were persistent (**Figure 1E**) and defined by fewer than 20 unique DARs, suggesting that 74% of the cells during development of ENZ-resistance share an overall similar chromatin accessibility profile. We then assessed changes in cluster chromatin DARs between the parental LNCaP, LNCaP-ENZ48, and in RES-A and RES-B. The RES-A specific cluster 6 (**Figure 1E**; 9% of the cells) had DARs at the *MYC* 5’UTR, *TP53* locus, and proximal to genes involved in steroidogenesis and cholesterol synthesis, consistent with activation of MYC and tendency to AR signaling re-activation. Conversely, the RES-B specific cluster 7 (10% of the cells) had DARs proximal to *PTPN12, CD177, and ADCY7* genes with functions in capturing external signaling and energy sources through membrane associated processes.

In line with studies on PC cell lines cultured for an extended time without androgens, which tend to display neuroendocrine-like phenotypes (Braadland et al., 2019; Fraser et al., 2019), the largest fold changes in chromatin accessibility based on average signal from all cells showed enrichment for neural system and neurite development processes between the parental (LNCaP-ENZ48 or DMSO) and resistant cells (RES-A or RES-B) (p < 0.001 in RES-A and p = 0.001 in RES-B). Using Gene Set Variation Analysis (GSVA) for gene expression scoring (see **Methods)**, we found elevated expression of NEPC-derived signatures among upregulated genes (Braadland et al., 2019; Tsai et al., 2017) in RES-A and RES-B cells (particularly *EZH2, AURKA, STMN1, DNMT1,* and *CDC25B*), as well as increased expression of NEPC-downregulated genes in initial clusters (**Figure S1G**).

Interestingly, in the same cell lines, deconvolution of bulk RNA-seq data with NEPC signatures showed high NEPC signal in RES-A cells only (**Figure S1H**), reflecting the lower resolution of bulk RNA-seq with respect to identifying signals from a small number of cells. Overall, these data show extensive chromatin reprogramming during emergence of resistance to AR-targeting agents.

Each sample comparison is indicated using colored dots within the plot and the *t*-test p-value is shown with asterisks (*** p-value < 0.001). **(C)** Venn diagram of shared and unique chromatin regions in LNCaP parental, LNCaP-ENZ48, RES-A, and RES-B according to bulk analysis of scATAC-seq. **(D)** UMAP scATAC-seq clustering visualization of LNCaP parental, LNCaP-ENZ48, RES-A, and RES-B. **(E)** Proportions of cells in scATAC-seq clusters.

Clusters are colored according to cluster type: initial (present in prevalence in LNCaP parental and LNCaP-ENZ48), ENZ-induced (present in prevalence in RES-A or RES-B), or persistent (present in similar proportions in all samples). See also Figure S1.

### Enzalutamide resistance reconfigures availability of TF binding DNA motifs in the chromatin

Chromatin accessibility determines transcriptional output by exposing a footprint of TF DNA motifs. As ENZ resistance was associated with increased chromatin opening, we hypothesized that this would change the footprint of TF DNA motifs exposed. To this end, we first used AR and MYC binding site maps in LNCaP cells (Barfeld et al., 2017) and explored the relationship between open chromatin sites according to the bulk FAIRE-seq data in RES-A cells. Using read distribution analysis, we observed a significant increase in open chromatin at MYC binding sites in ENZ-resistant cells (p < 0.001 in castrate conditions and with androgens, *t*-test) (**Figure 2A, Figure S2A**), as well as a reduction of open chromatin at AR binding sites (p < 0.001 in castrate conditions and with androgens, *t*-test) (**Figure 2B, Figure S2B**). These findings suggest that chromatin dysregulation in ENZ-resistance is associated with reconfiguration of AR and MYC chromatin binding, consistent with previously reported increased MYC and reduced AR transcriptional activity in these cells (Handle et al., 2019).

**Figure 2.**
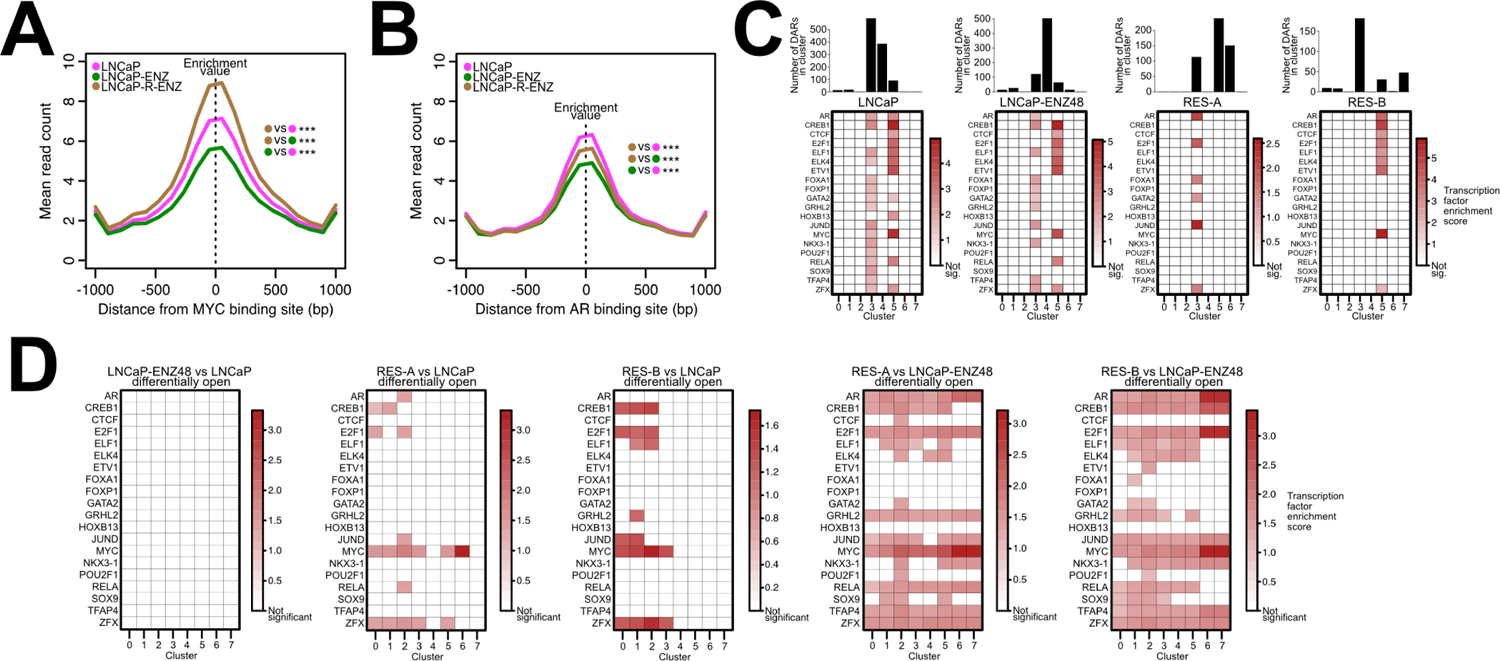
Contribution of enzalutamide treatment-mediated chromatin reprogramming to transcription factor DNA motif footprint. (A-B) Mean FAIRE-seq read count distribution in androgen-deprived conditions within a 2kb interval around MYC binding sites (A) and AR binding sites (B) in LNCaP cells. (C) Prostate cancer-associated transcription factor (TF) motif enrichment in open differentially accessible regions (DARs) for each scATAC-seq sample. Enrichments with a Benjamini-Hochberg method adjusted hypergeometric test p-value < 0.05 are shown in colors, while non-significant enrichment are shown in white. The barplots above the matrices indicate the number of open DARs found for each cluster in each sample. (D) TF motif enrichments in open DARs observed comparing the indicated conditions. Enrichments with a Benjamini-Hochberg method adjusted hypergeometric test p-value < 0.05 are shown in colors, while non-significant enrichment are shown in white. See also Figure S2.

To resolve how chromatin reprogramming affects TF DNA motif exposure at the single-cell level, we performed TF motif enrichment analysis on the marker DARs characterizing the scATAC-seq cell clusters in each sample (**Figure 2C**). This analysis confirmed enrichment of motifs for several PC-associated TFs such as AR and MYC, as well as GATA2, HOXB13, and others in persistent clusters 3 and 5 in parental and LNCaP-ENZ48 (**Figure 2C**). In stark contrast, only cluster 3 in RES-A and cluster 5 in RES-B showed enrichment of a subset of the same TFs motifs (**Figure 2C**), suggesting divergent TF dependencies for AR, CREB1, CTCF, E2F1, ETS-like, FOXA1, GATA2, JUND, MYC, and ZFX in these cell clusters in ENZ-resistant cells.

Between pairs of samples, DARs were predominantly opening in cluster 3 compared to all other clusters (8% vs 4% DARs differentially opening, *p* < 0.001, chi-square test) and predominantly closing in cluster 4 (11% vs 8% DARs differentially closing, *p* < 0.001, chi-square test) (**Figure S2C**). We performed selective TFs motif enrichment analysis in DARs opened (**Figure 2D**) and closed (**Figure S2D**) between pairs of samples (see **Methods**). While we observed no enrichments after short-term ENZ-treatment (LNCaP-ENZ48 vs parental; **Figure 2D**), comparing open DARs in RES-A or RES-B vs LNCaP parental retrieved distinct sets of TFs, with MYC and ESR1 being the most common across all clusters in RES-A and RES-B, respectively (**Figure 2D**). Similarly, comparing open DARs in RES-A or RES-B vs LNCaP-ENZ48 showed enrichment of most of the PCa-related TF motifs tested in most clusters (**Figure 2D**), and to an even greater extent when considering closing DARs between sample conditions (**Figure S2D**).

These analyses demonstrate that ENZ-resistance is associated with reconfiguration of TF DNA motif footprints, which is consistent with alteration of TFs activity (Hankey et al., 2020). DNA motifs for pioneer factors such as FOXA1 and GATA2 (Wang et al., 2007; Zhao et al., 2016) were not enriched in differentially open regions but were enriched in all clusters in the differentially closed regions (**Figure S2D**), possibly indicating a loss of activity at these sites (Hankey et al., 2020; Sahu et al., 2011). This is consistent with distinct chromatin-related mechanisms of ENZ-resistance observed in other bulk models (Zhang et al., 2020), meaning chromatin structure reprogramming in ENZ resistance might therefore be associated with TF binding reprogramming in a small subset of PC cells.

### Transcriptional patterns of enzalutamide resistance are induced by divergent chromatin reprogramming

To study transcriptional patterns in relation to reconfiguration of chromatin structure at the single-cell level, we additionally performed scRNA-seq in the LNCaP parental, RES-A and B models (**Figure 1A**). Visualizing clusters of cell subpopulations for the four samples (**Figure 3A**) showed 7 persistent, 3 ENZ-induced, and 3 initial cell clusters (**Figure 3B**) defined by sets of marker differentially expressed genes (DEGs; between 17 and 283 DEGs in the 13 clusters). To confirm that these cell subpopulations would be relevant in other independent models of ENZ-resistance, we validated the scRNA-seq clustering by using cluster label transfer (see **Methods**) to three additional scRNA-seq datasets (**Figure 1A**). We queried for matching cell populations in an additional LNCaP parental sample, LNCaP ENZ-treated for 1 week (LNCaP-ENZ168), and an independent ENZ-resistant, LNCaP-derived cell line (RES-C). Transferring scRNA-seq cluster labels confirmed the presence of initial clusters 4, 6, and 10 in RES-C (**Figure S3A**). We could find 79% of RES-B-specific ENZ-induced cluster 3 in LNCaP-ENZ168, suggesting that one week of ENZ treatment is sufficient to give rise to this cluster prior to the development of resistance (**Figure S3B**). Moreover, 17% of RES-C cells are coinciding with RES-B-specific ENZ-induced cluster 3, indicating that RES-B and RES-C lines share cells with similar characteristics. Most importantly, we could retrieve persistent subpopulations of cells in the alternative LNCaP-parental sample (4%; **Figure S3C**), in LNCaP-ENZ168 (13%), and in RES-C (31%), suggesting that these persistent cells are consistently found during emergence of ENZ-resistance.

**Figure 3.**
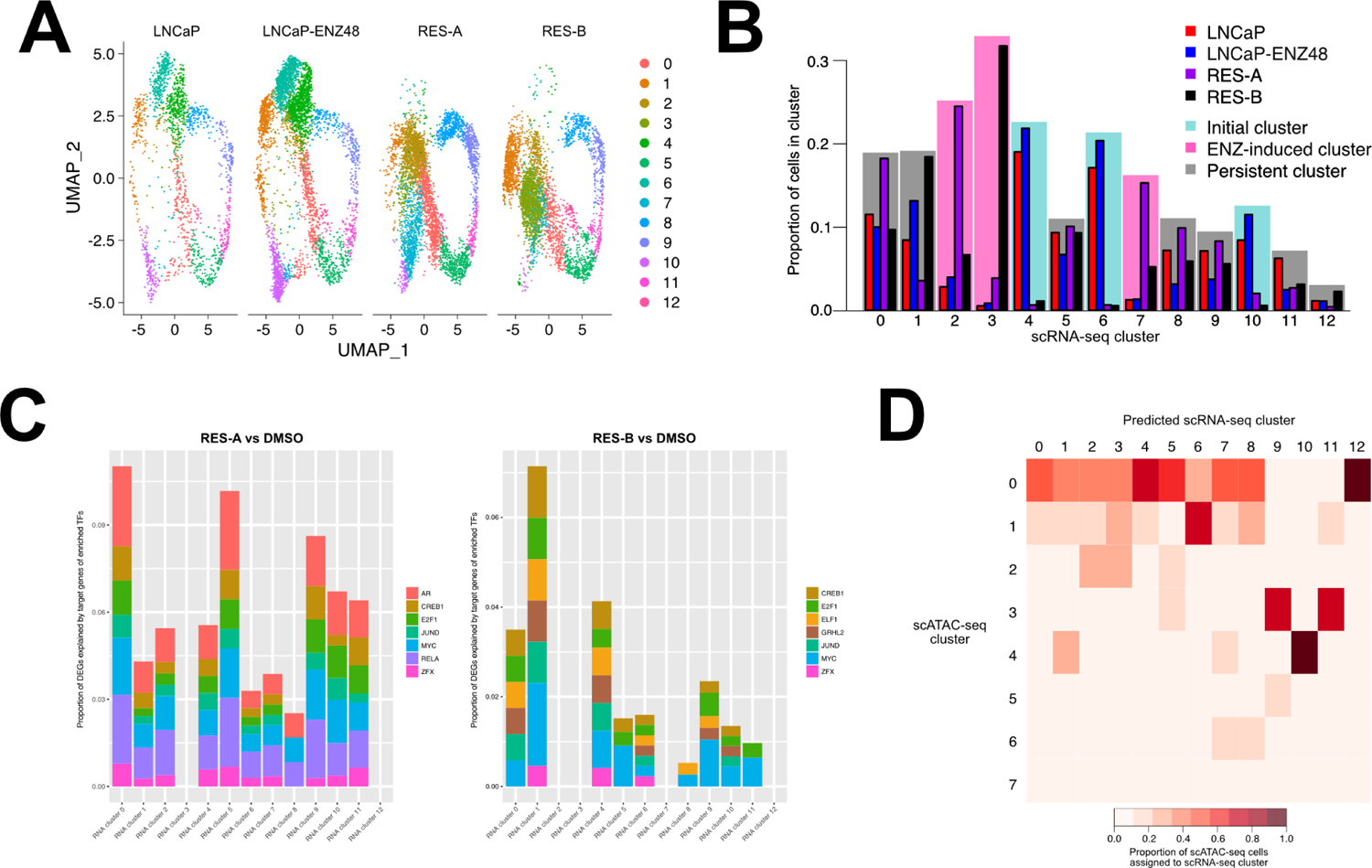
Chromatin states of enzalutamide resistance can result in multiple transcriptional programs. **(A)** UMAP clustering visualization of single-cell RNA sequencing (scRNA-seq) of LNCaP parental, LNCaP-ENZ48, RES-A, and RES-B. **(B)** Proportions of cells in clusters identified from scRNA-seq. Clusters are colored according to cluster type: initial (present in prevalence in LNCaP parental and LNCaP-ENZ48), ENZ-induced (present in prevalence in RES-A or RES-B), or persistent (present in similar proportions in all samples). **(C)** Proportion of differentially expressed genes in each scRNA-seq cluster for the indicated sample comparisons that is composed of enriched transcription factor (TF) target genes. The contributions of enriched TFs identified in the scATAC-seq are shown as a stacked barplot. **(D)** Identification of matching cell clusters between the scRNA and scATAC-seq data visualized as heatmap. The heatmap shows the proportions of scATAC-seq cells across all sample conditions assigned to each scRNA-seq cluster as part of the label transfer process. The proportions were calculated for each scRNA-seq cluster, with the total as the number of cells from the scATAC-seq that could be confidently assigned to a scRNA-seq cluster (confidence score > 0.4). See also Figure S3.

We then sought to determine whether the observed scRNA-seq clusters (**Figure 3A**) could be the result of enriched TFs binding activity in alternative open DARs. Using annotated databases, we queried the transcriptional targets of the enriched TFs in the open DARs when comparing RES-A or B to the parental LNCaP (**Figure 2D**) in the matching scRNA-seq samples (see **Methods**). Chromatin remodeling affected TF activity and consequently DEGs in the scRNA-seq for up to a maximum of 11% in cluster 0 in RES-A and 7.1% in cluster 1 in RES-B (**Figure 3C**). While target DEGs for TFs such as MYC, JUND, and E2F where promiscuously found in most clusters in both RES-A and B, other target DEGs for TFs such as AR, RELA (a NFkB subunit), and GRHL2 seemed more specific to RES-A or B, consistent with proposed stoichiometric models of TFs chromatin binding (Klemm et al., 2019). This analysis confirmed that alternative open DARs in ENZ-resistance can activate divergent transcriptional programs.

Next, we retraced scRNA-seq clusters to their scATAC-seq clusters. We took advantage once again of a label transfer approach to identify matching scRNA- and scATAC-seq cell states in the same sample conditions (see **Methods**). In this process, we assigned cell labels within the scRNA-seq to the scATAC-seq clusters, or vice versa (**Figure S3D**). We found that a chromatin state can correspond to multiple transcriptional states (96% in scATAC-RNA vs 48% in scRNA-ATAC of cells assigned on average across all samples, *p* < 0.001, chi-square test). Querying the integrated scRNA-seq clusters (**Figure 3A-B**) from the scATAC-seq data, we could find matching cell states in the scATAC-seq for scRNA clusters 9, 10, and 11 (**Figure S3E**). Across the sample conditions, 95% of the cells projected to belong to scRNA-seq cluster 10 belonged to scATAC-seq cluster 4, while 72% of cells projected to belong to scRNA-seq cluster 9 or 11 belonged to scATAC-seq cluster 3 (**Figure 3D**). Taken together, these data show that transcriptional configuration of ENZ-resistant cells, especially cells persisting during treatment, emerges from processes driven partially by chromatin structure and TF-mediated transcriptional reprogramming. These processes affect a number of important regulators of cell fate, consistent with lineage commitment recently observed in tissue development (Ma et al., 2020a).

### Prostate cancer cell subpopulations with features of stemness precede enzalutamide resistance

Cell cycle phase can be a strong determinant of the integrative clustering of our scRNA-seq data. Accordingly, we found that persistent clusters 8, 9, and 11 scored highly for S and G2/M phase related genes using cell cycle scoring in Seurat (see **Methods**) (**Figure 4A**), suggesting that cells in these clusters are more actively cycling and proliferative. However, we found that cells in clusters 9 and 11 were characterized not only by cell cycle genes, but also by expression of genes involved in chromatin remodeling and organization (CTCF, LAMINB, ATAD2), increased cell cycle turnover and stemness (FOXM1, (Ketola et al., 2017), and DNA repair (BRCA2, FANCI, RAD51C, POLQ) (**Figure 4B**). Clusters 5 and 11 showed high expression of a gene set, which we named “Stem-Like”, composed of stemness-related genes mainly from Horning *et al (Horning et al., 2018)* (**Figure 4C**). Karthaus *et al* recently identified activated luminal prostate cells able to regenerate the epithelium following castration (Karthaus et al., 2020). We extracted (see **Methods**) the gene expression profile associated with these prostate luminal cells, and used it to score each scRNA-seq cluster. We found cluster 10, an initial cluster, to score highly for this gene signature, which we renamed PROSGenesis (**Figure 4D**).

**Figure 4.**
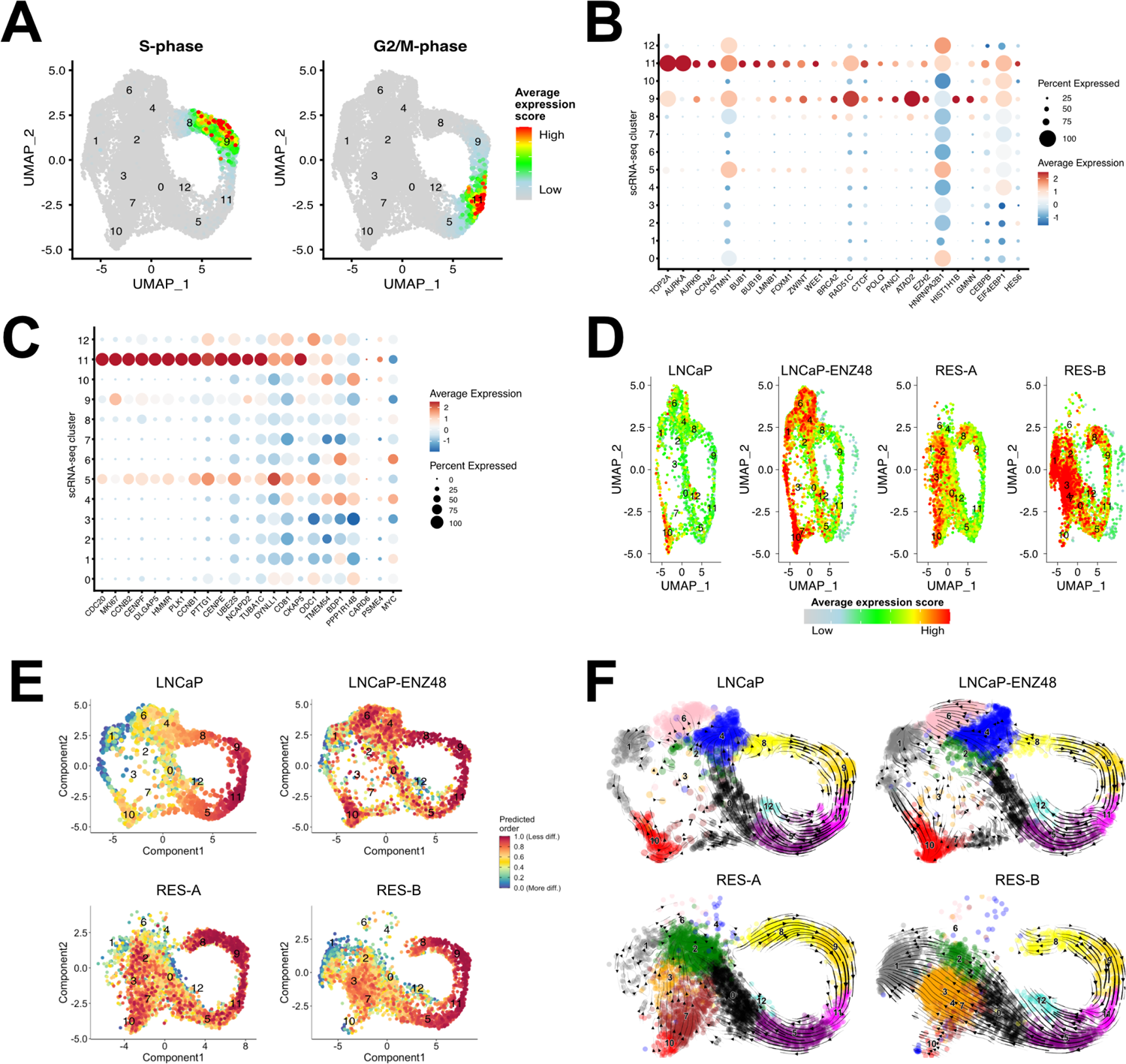
Transcriptional states of stemness in enzalutamide resistance. **(A)** Average expression of cell-cycle related genes (S and G2/M phases) in cells from the scRNA-seq data. **(B-C)** Dot plot of average gene expression of the (**B**) indicated genes and of the (**C**) genes within the Stem-Like signature in each scRNA-seq cluster. The size of the dot reflects the percentage of cells in the cluster that express each gene. **(D)** UMAP visualization showing the average expression score of each cell for the genes in the PROSGenesis gene signature derived from Karthaus *et al* (Karthaus et al., 2020). **(E)** Predicted differentiation states of cells in the four LNCaP scRNA-seq samples. Each cell is a dot colored according to its differentiation state. The scRNA-seq clusters are labeled with numbers. **(F)** RNA velocities based on scRNA-seq depicted as streamlines. Clusters are shown in different colors and are numbered. See also Figure S4.

We then set out to reconstruct the trajectories of how these clusters of interest were generated during the development of ENZ-resistance. Based on cytoTRACE (Gulati et al., 2020), cells in clusters 10 and 11 were the least differentiated across most of the sample conditions (**Figure 4E**), suggesting that the other cells could derive from cells in these clusters. RNA velocity analysis estimated cluster 10 as a precursor of the enzalutamide-induced clusters (**Figure 4F**), concordant with a state derived from activated regenerative luminal prostate cells as previously suggested (Karthaus et al., 2020).

Cluster-specific differential velocity analysis in RES-A and RES-B revealed downregulation of many PC-related genes, such as ATAD2, as well as upregulation of genes such as UBE2T, PIAS2, PFKFB4, and EGFR **(Figure S4A-B)**. ATAD2 and UBE2T were otherwise upregulated in persistent clusters 8, 9, and 11 (**Figure S4B**), suggesting additional transcriptional reprogramming in ENZ-induced clusters.

These analyses point at two distinct subpopulations of PC cells which precede ENZ resistance: one persistent cell cluster (cluster 11) matching “Stem-Like” and one initial cluster (cluster 10) matching PROSGenesis, a signature derived from tissue regeneration (Karthaus et al., 2020). Collectively, our data suggest that there exists a small number of PC cells within the bulk with stem-like and regenerative potential.

### Model-based characterization of gene signatures in prostate cancer bulk RNA sequencing

The use of molecular gene classifiers or signature scores is an attractive strategy to select cancer patients for treatment (Doultsinos and Mills, 2021; Eggener et al., 2020). Effectively, gene expression deconvolution methods are based on the use of gene signatures to assign cell tissue types from bulk RNA-seq (Wang et al., 2018). According to an unbiased enrichment and differential expression analysis of hallmark gene sets (see **Methods**), most of the persistent clusters and cluster 10 also showed significant enrichment of E2F targets, G2M checkpoint, and MYC target genes (**Figure S4C**). These data are largely concordant with the bulk RNA-seq data on the same cells in our previous study (Handle et al., 2019), reflecting the fact that signals from subpopulations of cells can be retrieved in bulk RNA-seq data. Differential expression within clusters (**Figure S4D-F**) further revealed that oxidative phosphorylation was immediately upregulated in LNCaP-ENZ48, and this process is maintained highly selectively in RES-A but not in RES-B. Moreover, genes regulated by activated mTORC1 signaling were consistently upregulated in most of the clusters as ENZ resistance develops (**Figure S4D-F**), in agreement with previous reports showing its activation during ENZ treatment in patients (Ma et al., 2020a).

We therefore used a collection of signatures derived from the scRNA-seq analysis to describe features of the same cells in bulk RNA-seq datasets. In addition to Stem-Like and PROSGenesis, we included (1) NEPC markers (**Figure S1G**), (2) a BRCAness gene signature (Li et al., 2017b), as RES-A and RES-B maintain sensitivity to PARP inhibition (Handle et al., 2019) and the persistent cluster 11 is characterized by markers of DNA repair (**Figure 4B**), (3) gene sets as proxies of AR signaling activation (He et al., 2018), including activation of AR splice variants (AR-Vs), (4) the DEGs defining our scRNA-seq clusters, and (5) gene sets for mTORC1 signaling and MYC targets (**Figure S4C-F**).

In the bulk, the ENZ-induced DEGs selectively appeared in the RES-B cells (**Figure 5A**). Similarly, the persistent clusters were associated with the Stem-Like signature only in RES-A and RES-B (**Figure 5A**). On the other hand, the PROSGenesis signature was elevated only in RES-B (**Figure 5A**). The NEPC features in RES-A were associated with MYC activation (**Figure 5A**). Consistent with both lines still being responsive to PARP inhibitors (Handle et al., 2019), we found samples from RES-A and RES-B to score highly for BRCAness (**Figure 5A**), which is known to downregulate DNA repair machinery (Li et al., 2017). BRCAness was associated with the AR-V signature as previously shown (Kounatidou et al., 2019) (**Figure 5A**).

**Figure 5.**
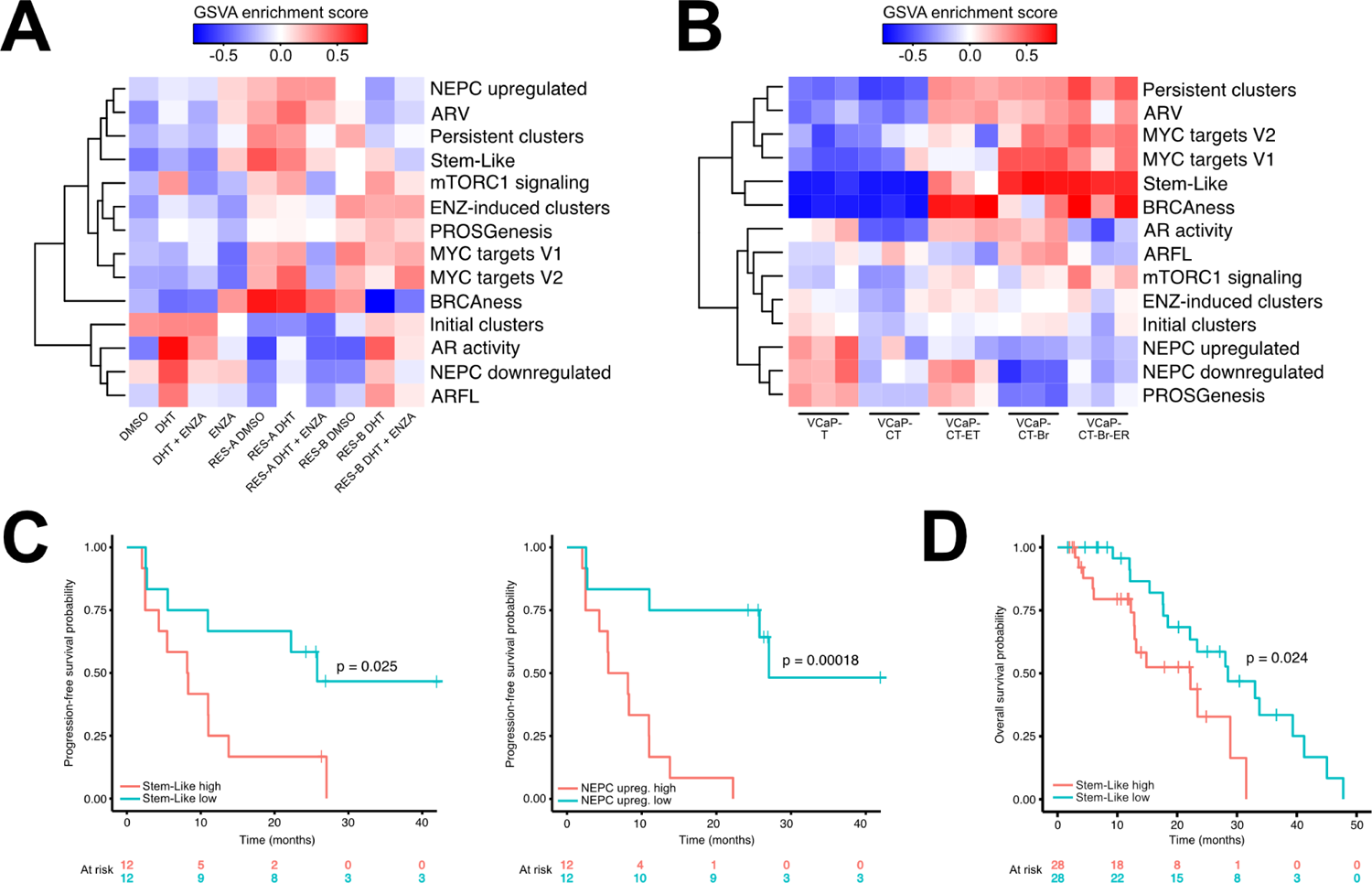
Gene signatures derived from single-cell RNA sequencing capture important features of prostate cancer models and stratify advanced PC patients. (A) Heatmap of gene signature GSVA enrichment scores in bulk RNA-sequencing of LNCaP treated with DHT or enzalutamide, and either sensitive or resistant to enzalutamide. (B) Heatmap of gene signature GSVA enrichment scores in bulk RNA-sequencing from VCaP subline derivatives VCaP-T (long term cultured with 10 uM testosterone), VCaP-CT (VCaP-T long term cultured with 0.1 nM testosterone), VCaP-CT-ET (VCaP-CT cultured long term with 10 µM enzalutamide), VCaP-CT-Br (VCaP-CT cultured long term with bicalutamide), and VCaP-CT-Br-ER (VCaP-CT-Br long term treated with enzalutamide upon reaching bicalutamide insensitivity). (C) Kaplan-Meier curves for Alumkal *et al* patients stratified into two groups based on median GSVA score for Stem-Like and NEPC upregulated gene signatures. Log-rank p-value is indicated above the curves. (D) Kaplan-Meier curve for abiraterone and enzalutamide-naive patients from SU2C (bone metastasis samples excluded) stratified into two groups based on median GSVA score for the Stem-Like gene signature. Log-rank p-value is shown above the curve. See also Figure S5.

To confirm the properties of different signatures, we used VCaP cells to develop an independent model of resistance to AR signaling-targeted treatments including ADT, bicalutamide, ENZ, and bicalutamide/ENZ multi-resistant sublines, and performed bulk RNA-seq (**Figure 1A**). These VCaP-based sublines did not show NE features (**Figure 5B**). Only ENZ-resistant VCaP cells scored highly for the ENZ-induced DEGs, confirming the specificity of this signature to ENZ treatment and resistance. Parental and ENZ-resistant VCaP cells scored highly for the PROSGenesis signature, while the scores of the persistent, Stem-Like, mTORC1 signaling, and MYC targets signatures scored high selectively in resistant VCaP sublines (**Figure 5B**). This suggests a convergent mechanism of resistance to these agents in this independent model.

Next, we scored xenografts of AR^+^/NE^-^, AR^-^/NE^+^, or AR^-^/NE^-^ CRPC and NEPC tumors resistant to ENZ (Labrecque et al., 2019; Lam et al., 2020) with the same signature sets (**Figure S5A**). AR^+^/NE^-^ xenograft samples clustered into two separate clusters. AR^-^ tumors clustered together with a series of AR^+^/NE^-^ tumors due to low mTORC and MYC signaling, while one cluster of AR^+^/NE^-^ scored highly for all of the gene sets except for markers upregulated in NEPC “NEPC upregulated”. Interestingly the PROSGenesis signature, along with initial clusters and ENZ-induced clusters, scored particularly high in AR^+^ tumors while the Stem-Like signature, along with the persistent clusters, scored high in both AR^+^/NE^-^ and AR^-^/NE^+^ tumors (**Figure S5A**), suggesting that the two signatures capture different tumor biologies. In a transcriptome dataset based on an independent xenograft model (King et al., 2017), we found ENZ resistance to be uniquely associated with higher AR activity, higher expression of MYC target genes, PROSGenesis high score, and high expression of ENZ-induced cluster gene sets (**Figure S5B**). This data suggest that the Stem-Like status is independent of the AR status and that persistent cells might mediate the development of both AR positive CRPCs and negative NEPCs. Collectively, the persistent, initial, PROSGenesis, and Stem-Like derived gene signatures show potential for identifying aggressive regenerative features of PC from bulk RNA-seq.

### Transcriptional signal deconvolution identifies treatment persistent cells and prognostic gene signatures in prostate cancer patients

We then hypothesized that we could systematically use gene signatures as a proxy for the presence of PC cells with different transcriptional features in clinical settings. We first interrogated clinical data of CRPC patients treated with ENZ reported in Alumkal et al. (Alumkal et al., 2020). The patients aggregated into two clusters based on our complete signature set (**Figure S5C**), but patients in neither cluster had significantly shorter progression-free survival (PFS; p > 0.05, log-rank test). Utilizing a stepwise variable selection process we identified five significant signatures (NEPC upregulated, PROSGenesis, MYC targets, AR activity, and ARV) that are able to identify patients with significantly shorter PFS (**Figure S5D**). Moreover, PFS analysis of individual gene signatures revealed association with shorter time to progression for patients scoring high for the Stem-Like signature (p = 0.025, log-rank test) or for genes upregulated in NEPC (p < 0.001, log-rank test) (**Figure 5C**), while patients with longer PFS scored highly for PROSGenesis (p = 0.021, log-rank test) and for the initial cluster signature (p = 0.018, log-rank test) (**Figure S5E**).

None of the cluster marker gene sets showed a significant difference between Stand Up To Cancer (SU2C) CRPC abiraterone/ENZ-naive and abiraterone/ENZ-exposed patients (Abida et al., 2019) according to their latest treatment regime, suggesting that differences between the tumors based on the signatures may be difficult to retrieve using bulk sequencing from heavily pre-treated patients. Despite the challenges of applying single-cell derived signatures to bulk data however, Stem-Like was still significantly associated with poor overall survival in these patients (**Figure 5D**), supporting the potential significant activity of the persistent cells in this group of patients. Similarly, we could not stratify patients that developed resistance to ENZ in the SU2C West Coast DT Quigley *et al* dataset (**Figure S5F**) (Quigley et al., 2018), although in this case, ENZ-sensitive patients had higher expression of PROSGenesis (p = 0.024, Wilcoxon rank-sum test) (**Figure S5G**).

These data show that the Stem-Like signature associated with persistent cells (cluster 11) from our single-cell analysis of ENZ resistance is a consistent classifier with the potential of stratifying patients for response to second line AR-targeted treatments (**Table 1**). The findings also support the presence of a small number of cells within the tumor that score high for Stem-Like and PROSGenesis, with the potential ability to regenerate the bulk during treatment (Zahir et al., 2020).

**Table 1.** Summary table of gene signature GSVA score associations with PFS or OS in the clinical datasets. Only gene signatures significantly associated with progression-free survival (PFS) or overall survival (OS) in one or more datasets are shown. Good indicates a higher score for the signature (a score higher than the median) is associated with better survival outcome, while poor indicates that a higher signature score (a score higher than the median) is associated with worse survival outcome. Log-rank p-values are shown with asterisks (* p-value < 0.05, ** p-value < 0.01, *** p-value < 0.001). For each dataset, the header indicates the number of samples included, along with other qualifying information of the dataset. We used abiraterone (ABI)/ENZ naive patients from the Stand Up To Cancer (SU2C) CRPC dataset.

| Gene signature | Alumkal *et al*<br>(PFS)<br>(n = 24) | SU2C CRPC<br>(OS)(capture,<br>ABI/ENZ<br>naive)<br>(n = 56) | Quigley *et al* (OS)<br>(n = 96) | TCGA-<br>PRAD<br>(PFS)<br>(n = 271) | ICGC- EOPC<br>(PFS)<br>(n = 85) |
| --- | --- | --- | --- | --- | --- |
| Initial clusters | Good* |  |  |  |  |
| PROSGenesis | Good* |  |  | Good** |  |
| Cluster 10 | Good* |  |  | Good** |  |
| Cluster 4 | Good** |  |  |  |  |
| ENZ-induced<br>clusters |  |  |  | Good*** |  |
| Cluster 3 |  |  |  | Good*** |  |
| Persistent clusters |  |  |  | Poor** |  |
| Stem-Like | Poor* | Poor* |  | Poor* |  |
| Cluster 11 | Poor* |  |  | Poor*** |  |
| Cluster 9 |  |  |  | Poor*** |  |
| Cluster 5 |  |  |  | Poor*** |  |
| Cluster 0 |  |  |  | Poor** |  |
| Cluster 1 |  |  |  | Good*** |  |
| Cluster 8 |  |  |  | Poor** |  |
| AR activity |  |  |  | Good* |  |
| ARFL |  |  |  | Good* |  |
| NEPC upregulated | Poor*** |  |  |  |  |
| NEPC<br>downregulated |  |  |  |  | Good* |
| BRCAness | Poor** |  |  | Poor** |  |

Using PC specimen tumor DNA, we recently showed the presence of subclones within the primary tumors that preserve the ability to expand and metastasize years after treatment and are found interlayered within different lesions of multifocal tumors (Woodcock et al., 2020). Similarly, a recent work studying lung cancer metastases found that metastatic capacity arises from pre-existing and heritable differences in gene expression (Quinn et al., 2021). Therefore, we hypothesized that the persistent cluster 11, Stem-Like, initial cluster 10, and PROSGenesis signatures could be able to capture signals from such types of pre-existing subclones with metastatic potential in primary untreated tumors.

To this end, we took advantage of a recently published scRNA-seq dataset on clinically relevant PCs specimens (**Figure S6A**) (Chen et al., 2021). We used GSVA score to highlight our 13 scRNA-seq clusters in 36424 cells from primary untreated PC specimens of 13 patients (**Figure S6B**). The analysis showed that our LNCaP model-derived cell clusters scored higher in luminal and basal/intermediate cells compared to fibroblasts (average score −0.07 vs −0.20, p = 0.047, *t*-test) (**Figure S6B**). Additionally, luminal cells had higher expression of genes associated with our initial scRNA-seq clusters compared to the basal/intermediate cells (average score 0.23 vs −0.12, p = 0.02, *t*-test) and compared to fibroblasts (average score 0.23 vs −0.39, p < 0.001, *t*-test).

We then scored the cells for expression of genes from the Stem-Like and PROSGenesis signatures, along with the associated clusters (11 and 10, respectively) and control signatures linked to AR activity (ARV, AR-FL, and AR activation), BRCAness, and NEPC (**Figure 6A**). We defined a high score for a gene signature to be above the 90th percentile. 48% percent of the cells that scored highly for the Stem-Like signature were luminal cells (**Figure 6B**). Cells scoring highly for the PROSGenesis signature were mostly basal/intermediate (78% of high scorers) (**Figure 6C**). Each single patient harbored on average 8% of cells scoring high for the Stem-Like signature (ranging from 2% in patient 173 to 23% in patient 156) and 8% of cells scoring high for the PROSGenesis signature (ranging from 0.9% in patient 153 to 33% in patient 172) (**Figure 6D**).

**Figure 6.**
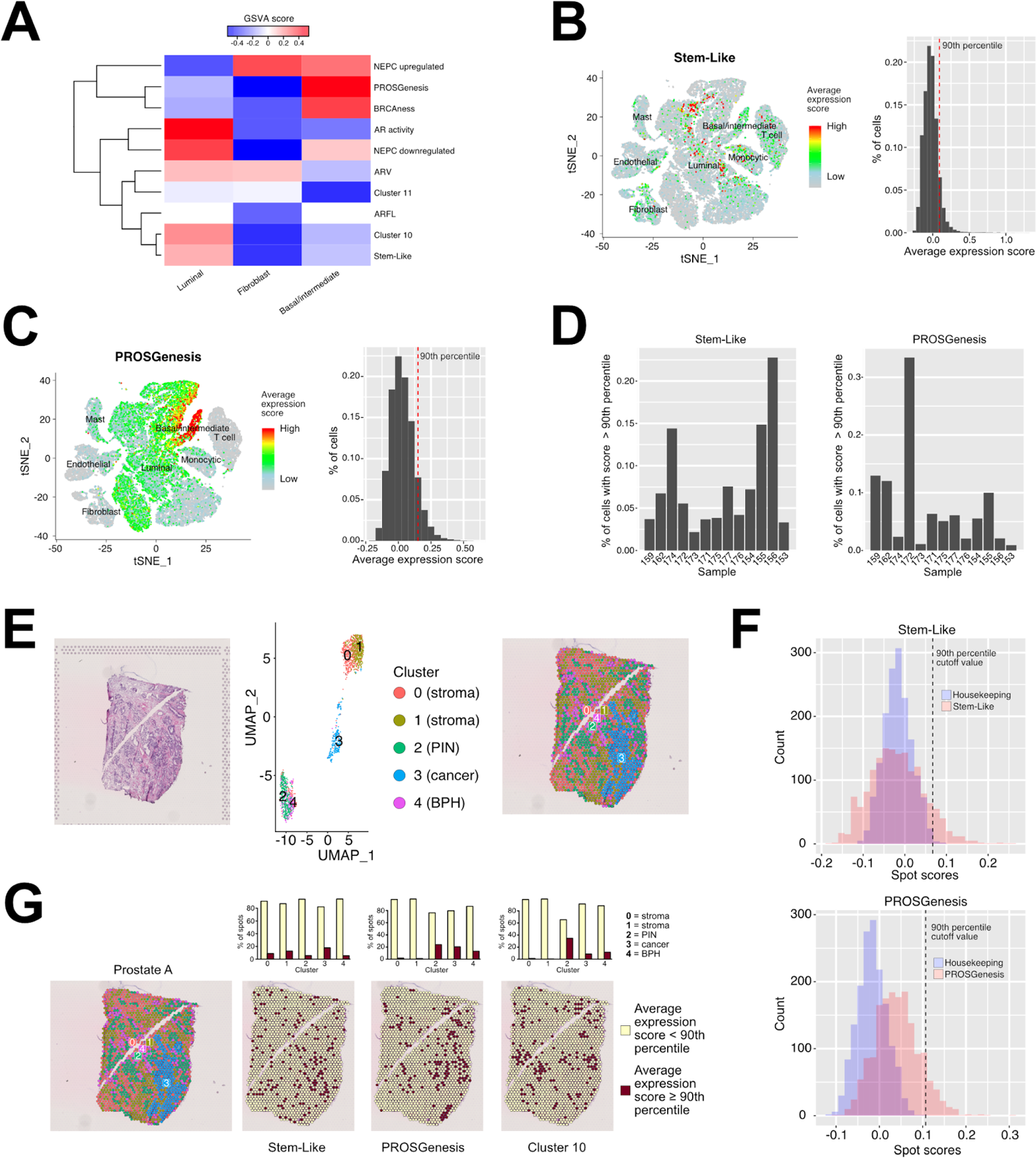
Transcriptional signal deconvolution identifies treatment persistent cells in prostate cancer. **(A)** GSVA enrichment scores from single-cell RNA-seq data for gene signatures in luminal, basal/intermediate, and fibroblast cells in specimens from 12 treatment-naive prostate cancer (PC) patients (Chen et al., 2021). GSVA enrichment scores were generated from the average expression profile of each cell type. See Figure S6 for the tSNE visualization of the cell types. **(B-C)** tSNE plot of PC cells colored according to their average expression of the genes in **(B)** the Stem-Like signature and in **(C)** the PROSGenesis signature. The adjacent histograms show the distribution of average expression scores in the cells, with a red dotted line denoting the 90th percentile of scores. **(D)** Percentage of cells scoring at or above the 90th percentile for the Stem-Like and PROSGenesis signatures belonging to each patient. **(E-G)** Spatial transcriptomics (ST) from a prostate cancer tissue section, Prostate A. **(E)** H&E staining of the tissue section (left most panel), UMAP visualization (central panel) of the clusters of the spots on the ST slide (right most panel). Each cluster is also labeled according to its histological tissue type, with clusters 0 and 1 corresponding to stroma, cluster 2 corresponding to prostatic intraepithelial neoplasia (PIN), cluster 3 corresponding to the prostate adenocarcinoma, and 4 corresponding to benign prostatic hyperplasia (BPH). **(F)** Sensitivity analysis of Stem-Like and PROSGenesis signatures scores in ST against the score distributions of control housekeeping gene signatures (see **Methods**). **(G)** The leftmost panel shows the ST UMAP clusters of spots overlaid on the H&E slide. Each spot was scored according to its expression of genes in the Stem-Like, PROSGenesis, and cluster 10 signatures. For each signature, spots scoring at or above the 90th percentile (“high”) are colored in red, while spots scoring below the 90th percentile (“low”) are colored in yellow. The barplots indicate the percentage of spots in each cluster scoring high or low for each signature. See also Figure S6.

To reconcile the presence of these cells and their relative histopathological position, we assessed gene expression within 2 sections of primary untreated PC with spatial transcriptomics (see **Methods**). We reconstructed the gene expression signal from stromal and epithelial components in an average of 1682 spots per sample using clustering analysis and annotated the tissue architecture in 5 clusters of different morphologies of benign stromal tissue (BT), benign prostatic hyperplasia (BPH), prostate intraepithelial neoplasia (PIN), and adenocarcinoma (PC-AC) (**Figure 6E, Figure S6C**). PROSGenesis and Stem-Like signatures, as well as the companion model-derived cluster 10 signature, showed high expression scores compared to scores from a housekeeping gene signature (**Figure 6F, Figure S6D**). We compared the score distributions of our signatures to the housekeeping gene set score distributions, and determined the 90th percentile as a score cutoff for high expression by allowing for 5% false positives (see **Methods**). Spots with high signal were found interspersed in all 5 clusters (**Figure 6G, Figure S6E**). However, we observed enrichment for spots scoring highly for the Stem-Like signature in the PT-AC cluster compared to benign tissue (18% vs 11% high scoring spots, p = 0.005, chi-square test). Spots scoring highly for PROSGenesis were enriched in the PIN and PT-AC clusters compared to benign tissue (p < 0.001 in both cases, chi-square test), while spots scoring highly for cluster 10 were enriched in the PIN cluster compared to all other tissue regions (p < 0.001 for each comparison, chi-square test) (**Figure 6G**). Taken together these data suggest the presence of treatment-persistent cells interspersed within the primary untreated prostate tissue of PC patients with high metastatic potential.

We finally verified whether we could predict recurrence in primary PC patients using the signal derived from the signatures in these cells. We interrogated legacy primary tumor TCGA PRAD (https://www.cancer.gov/tcga) (**Figure 7A**) and early onset PC (EOPC) ICGC (Gerhauser et al., 2018) gene expression data (**Figure S7A**) for our gene signatures of interest. Using all signatures for clustering the TCGA PRAD cohort separated 54% of Gleason score (GS)-7 and 15% of GS-8+ patients which would not benefit from additional treatment, as they had relatively good prognosis (**Figure 7B**). A similar trend was also observed in the ICGC cohort (**Figure S7B**). ENZ-induced (**Figure 7C**), PROSGenesis (**Figure 7D**), Stem-Like (**Figure 7E**), and persistent (**Figure 7F**) gene signatures were the most significant (p < 0.05, log-rank test) contributors to cluster separation in the TCGA cohort, while NEPC downregulated genes were the major determinant in the ICGC cohort (**Figure S7C**), reflecting the different biology of these GS-7-enriched EOPCs. In the ICGC cohort, the persistent and PROSGenesis signatures significantly stratified GS-7 patients (p < 0.05, log-rank test) (**Figure S7D-E**), suggesting the ability of these signatures to further refine GS-based risk stratification in these patients and avoid overtreatment. High PROSGenesis score was associated with good prognosis together with the gene set from the initial cluster 10 (**Table 1**). In line with previous reports (Alumkal et al., 2020), signatures reflecting AR activity (AR activity and full length) in these tumors were consistently associated with longer time to progression in the TCGA cohort (**Figure 7G-H**), suggesting a better response to inhibition of AR signaling in AR driven tumors. Individually, 8 out 13 clusters-derived signatures showed association with PFS in the TCGA cohort (**Table 1**), pointing at the utility of these signatures in PC patient risk stratification.

**Figure 7.**
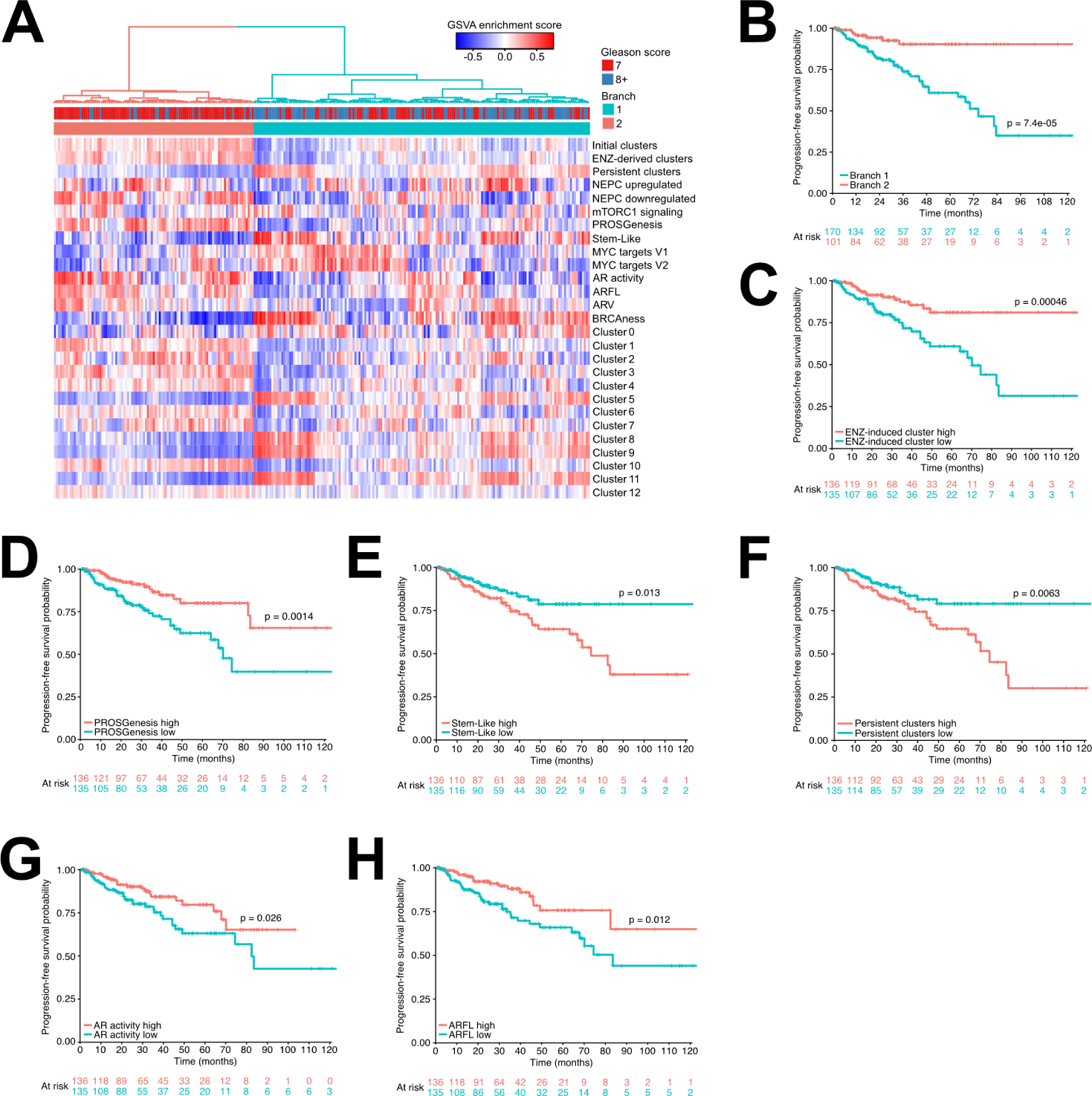
Transcriptional signal from persistent prostate cancer cells can be used to stratify untreated patients. **(A)** Heatmap of GSVA enrichment scores for all single-cell-derived gene signatures in the TCGA-PRAD cohort, including the marker gene sets for each scRNA-seq cluster. Hierarchical clustering of the GSVA scores was used to separate the samples into two groups, marked Branch 1 and Branch 2. **(B)** Kaplan–Meier survival curve for TCGA-PRAD patients stratified into two groups as indicated in Panel A. **(C-H)** Kaplan-Meier survival curves for TCGA-PRAD patients stratified into two groups based on median GSVA score for signatures of ENZ-induced cluster, PROSGenesis, Stem-Like, persistent cluster, AR activity, and ARFL. In each plot, the log-rank p-value is indicated above the plotted curves. See also Figure S7.

## Discussion

In this study we provide a molecular perspective of the evolution of resistance to AR targeted treatment at a single-cell level. Karthaus and colleagues recently found that luminal prostate cells that persist after ADT in a mouse model can contribute to tissue regeneration of the normal prostate epithelium by assuming stem-like transcriptional properties (Karthaus et al., 2020). Similar phenotypic features important for routing and rates of metastasis formation in lung cancer models have been found to be driven by differences in gene expression (Quinn et al., 2021). Similarly, here we find that during exposure to AR targeting agents, a small proportion of persistent cells remain transcriptionally unperturbed by the treatment, consistent with our recent findings of subclones with high metastatic potential interlayered within the primary untreated lesions (Woodcock et al., 2020).

We visualized these cells in primary untreated PC specimens and located them interspersed in cancerous regions of histopathologically relevant tumors as well as pre-lesions PIN and even in apparent benign tissue using spatial transcriptomics. Our data show evidence of a hierarchical model of emergence of resistance to enzalutamide (Maitland, 2021) in which treatment-persistent cells are able to regenerate the bulk of the resistant ones. We describe the properties of the persistent cells using RNA velocity and show different intermediate states in alternative trajectories of treatment resistance. This process is partially driven by chromatin remodeling, which is consistent with chromatin accessibility lineage-priming (Ma et al., 2020a; Martin et al., 2020). In PC, gain of function of bromodomain containing proteins such as BRD4 (Asangani et al., 2014; Urbanucci et al., 2017) and ATAD2 (Morozumi et al., 2016; Urbanucci et al., 2017), as well as loss of function of chromatin remodeler CHD (Zhang et al., 2020), have been shown to contribute to PC progression and lineage plasticity in therapy resistance. This process is likely accompanied by chromatin reprogramming (Braadland et al., 2019; Urbanucci et al., 2017; Uusi-Mäkelä et al.). While many groups focused on the effect of AR-targeted treatment on chromatin associated factors such as CREB5 (Hwang et al., 2019), or TFs such as GR (Arora et al., 2013) and AR (Yuan et al., 2019), in this study we found that exposure to AR-targeting agents increases the overall relaxation of the chromatin. The single-cell level analysis of the chromatin revealed that subpopulations of cells with different chromatin states lead to multiple transcriptional configurations, including those of persistent cells. Using different cell line models mimicking alternative trajectories of treatment-resistance, we infer that differential DNA motif exposure determined by chromatin structure may partially contribute to TF activity-mediated transcriptional reprogramming in the different cell subpopulations induced by exposure to enzalutamide. According to this analysis, specific subpopulations of PC cells are more subjected than others to TFs activity reprogramming. This is consistent with recent studies showing simultaneous detection of multiple transcription factors on single DNA molecules and TFs co-occupancy frequently occurring at sites of competition with nucleosomes (Strickfaden et al., 2020).

We show that treatment-persistent cells have high cell cycle turnover, compatible with high regenerative potential (Poli et al., 2018; Wang et al., 2020), and assume states of stemness from their transcriptional profiles. As these features have been associated with more aggressive tumors, we developed transcriptional signatures derived from two states in particular: one state derived in ADT-treated mice prostate cells by Karthaus et al. (Karthaus et al., 2020), which we renamed PROSGenesis and which tightly associated with initial and enzalutamide-induced clusters in our model of enzalutamide resistance, and one that we called “Stem-Like”, associated with persistent cells during evolution of enzalutamide resistance. Different signatures can capture different tumor types and inform treatment response. PROSGenesis, Stem-Like, and associated signatures derived from our enzalutamide-resistant models stratify ARSI exposed CRPC patients’ outcome. Moreover, we show that in primary PC patients undergoing ADT treatment, high signature scores in treatment-naive specimens are associated with short time to PFS (biochemical recurrence). Interestingly, in primary naive patients, high score for PROSGenesis is associated with longer lasting response to ADT, possibly due to a stronger contribution of AR activity in these tumors. Overall, we profile subpopulations of treatment-persistent cells with stem-like and regenerative properties that foster alternative evolutions of AR-targeted treatment resistant PCs.

## Methods

### Cell lines and culture

LNCaP cell lines were obtained from American Type Culture Collection (ATCC; LGC Standards) and authenticated periodically (HPA cultures or Eurofins). RES-A and RES-B cells were generated as described previously by prolonged exposure to the second-generation anti-androgens enzalutamide and RD-162 as described earlier (Handle et al., 2019). LNCaP parental (ATCC), RES-A, and RES-B cells were cultured in RPMI 1640 (Sigma R0883) supplemented with 10% FBS (Sigma F7524), 2 mM Alanyl-glutamine (Sigma G8541), 1 mM sodium pyruvate (Merck TMS-005-C), 2.5 g/L glucose (Sigma G8769), and 1x Antibiotic-Antimycotic (Gibco, 15240062) in a humidified 37°C incubator with 5% CO_2_.

RES-A and RES-B cells additionally received 10 µM enzalutamide (MedChemExpress HY-70002) with each cell splitting/feeding. For experimental treatments, ∼1×10^6^ cells were seeded into 5 cm culture plate dishes, and allowed to settle before exposure to 10 µM enzalutamide or DMSO vehicle control (0.1%) for 48 h or 168 h. The additional LNCaP cells (ATCC) and RES-C cells were cultured in a humidified CO2-incubator at 37°C in Gibco™ RPMI 1640 (1X) media (Thermo Fisher Scientific) supplemented with 10% FBS (Gibco standard FBS, Thermo Fisher Scientific), 2 mM L-Glutamine (Gibco®, Thermo Fisher Scientific), and a combination of 100 U/ml Penicillin and 100 μg/ml Streptomycin (Gibco® Pen Strep, Thermo Fisher Scientific). The enzalutamide resistant LNCaP RES-C cell line was generated by passaging of LNCaP cells with continuous treatment with 10 µM enzalutamide for 9 months and maintained in the same medium as LNCaP except for the supplementation with 10 µM enzalutamide.

### Generation of resistant VCaP subline derivatives and RNA-seq

Androgen-sensitive VCaP prostate cancer cell line (passage (p.) 15.) was a gift from Dr. Tapio Visakorpi, Tampere University, Finland. Cells were cultured in RPMI 1640 supplemented with 10 % DCC-FBS, 1 % L-glutamine, 1 % A/A, and 10 nM testosterone (T) for seven months to establish T-dependent subclone VCaP-T. VCaP-T cells were then cultured at low testosterone (0.1 nM) for 10 months to establish VCaP-CT, an androgen-independent cell line able to grow despite low testosterone. VCaP-CT were then cultured at 10 µM enzalutamide until the cells regained ability to grow despite enzalutamide, creating enzalutamide resistant cell line VCaP-CT-ET. Another cell line was created by incubating first VCaP-CT cells with bicalutamide and subsequently with enzalutamide upon reaching bicalutamide insensitivity. Ultimately these cells also gained the ability to grow despite enzalutamide, creating the multiresistant cell line VCaP-CT-Br-ER.

RNA sequencing was performed with Illumina HiSeq 3000. We sequenced 3 replicates, obtaining an average of 111 million paired-end reads per sample. Reads were aligned using STAR aligner version 2.5.4b and Ensembl reference genome GRCh38. Genewise read counts were quantified using featureCounts version 1.6.2 and Gencode annotations release 28.

### Single-cell samples preparation and sequencing

LNCaP parental, treated for 48 hours with enzalutamide or DMSO, RES-A, and RES-B cells were harvested with 0.05% Trypsin-EDTA (Sigma T3924). After neutralization with complete medium, centrifugation (300 x g for 5 min), and resuspension in PBS/0.5% BSA, the cells were filtered through a 35 µm Cell Strainer (Corning 352235) and a single-cell suspension of living cells was acquired through sorting on a FACS Aria II cell sorter. The cell concentration of the single-cell suspension was assessed with a Countess II FL Automated Cell Counter and ∼3 x 10^4^ cells were pelleted (300 x g for 5 min) for further processing for using the Chromium Single Cell 3’ Library, Gel Bead & Multiplex Kit, and Chip Kit (v3, 10x Genomics).

For the additional LNCaP parental and RES-C cells, 1 million cells were thawed in RPMI (Gibco) with 10% FBS (Gibco) and centrifuged at 300g for 5 min. The cells were then suspended in PBS with 0.04% BSA (Ambion) and filtered with Flowmni™ cell strainer (Bel-Art). Before loading, the cells’ viability and concentration was determined using Trypan blue with Cellometer Mini Automated Cell Counter (Nexcelom Bioscience). Chromium Single Cell 5’ RNA-seq was performed using the 10X Genomics Chromium technology, according to the Chromium Next GEM Single Cell V(D)J Reagent Kits v1.1 kit User guide CG000208 Rev D with loading concentration of 1000-200 cells/µl.

The LNCaP-ENZ168 single-cell RNA-seq sample was performed with Drop-seq (Macosko et al., 2015) using the Dolomite cell encapsulation system (Dolomite Bio). Cells were trypsinized with TrypLETM Express Enzyme (ThermoFisher Scientific, #12604021), spun down (5 min at 300xg) and washed with 0.1% BSA-PBS. After pelleting, the cells were resuspended in plain PBS and passed through a 40-micron filter. The number of viable cells was estimated with the use of trypan blue staining and Fuchs-Rosenthal hemocytometer chamber. The concentration of cells was brought down to 3×105 cells/mL in 0.1% BSA-PBS. For single-cell encapsulation, single-cell suspension, beads in lysis buffer and oil were connected with the loops and tubing to the Mitos P pumps and run through the glass microfluidic chip at the following flow rates: 100μL/min (Oil channel), 20μL/min (Bead channel); 350 mbar (Cell channel). Droplets were separated by centrifugation and beads counted with the use of Fuchs-Rosenthal hemocytometer chamber and up to 90000 beads were collected into one tube for Reverse Transcription reaction, exonuclease treatment, and amplification of cDNA library according to the original protocol (Macosko et al., 2015). Tagmentation of cDNA was performed with the Nextera XT DNA Library Preparation Kit (Illumina, #FC-131-1024). The PCR product was cleaned-up with AMPure XP beads, eluted in 10μL H2O and sequenced using Illumina HiSeq 2500 Rapid run.

For scATAC-seq, cell nuclei were isolated following the 10x Genomics Demonstrated Protocol for Single Cell ATAC Sequencing (CG000169-Rev C). Briefly, the cell suspension was washed once in PBS/0.04%BSA, and 2 x 10^5^ cells were pelleted (300 x g for 5 min), resuspended in 100 µl freshly prepared Lysis Buffer (10 mM Tris-HCl pH 7.4, 10 mM NaCl, 3 mM MgCl_2_, 0.1% Tween-20, 0.1% NP40 Substitute, 0.01% Digitonin, 1% BSA), and incubated on ice for 4 min (LNCaP parental cells), 6 min (RES-A), or 5 min (RES-B). The lysates were diluted with 1 ml wash buffer (10 mM Tris-HCl pH 7.4, 10 mM NaCl, 3 mM MgCl_2_, 0.1% Tween-20, 1% BSA), and the nuclei were pelleted (500 x g for 5 min) and resuspended in 30 µl 1x Nuclei Buffer (10x Genomics PN-2000153). Successful preparation of intact, isolated nuclei was confirmed through visual inspection in a phase-contrast microscopy, and nuclei concentration was assessed with a Countess II FL Automated Cell Counter, before proceeding immediately to processing for Single Cell ATAC sequencing using 10x Chromium, 10x Genomics library preparation and the Chromium Single Cell ATAC Reagent Kits (v1) User Guide (CG000168 Rev D).

Sequencing was performed on the Illumina NextSeq500 instrument at the genomics core facility at the Oslo University Hospital, while sequencing of the additional LNCaP parental and RES-C was performed with Novogene Company Limited, Cambridge, UK’s sequencing core facility was used with a PE150 NovaSeq sequencer, aiming at 50000 reads per cell.

For scRNA-seq, sequencing reads were processed into FASTQ format and single-cell feature counts using Cell Ranger v3.0.2 (Zheng et al., 2017). Similarly, Cell Ranger ATAC v1.1.0 (Satpathy et al., 2019) was used to process sequencing reads from scATAC-seq into FASTQ format and peak-barcode counts. The LNCaP-ENZ168 Drop-seq sample was pre-processed, aligned, and processed to a cell count matrix using Drop-seq tools v2.3.0 (Macosko et al., 2015). We utilized the human reference genome version GRCh38, along with NCBI RefSeq gene annotations for genome build GRCh38.p12.

### Formaldehyde-assisted isolation of regulatory elements (FAIRE) sequencing and analysis

FAIRE was performed on parental and LNCaP-ResA cells in biological triplicate according to the standard protocol (Simon et al., 2012). Prior to FAIRE-seq, cells were cultured for three days in RPMI medium supplemented with 5% DCC FBS and 10 µM enzalutamide was added only to the resistant cell line. Both sublines were then treated with DMSO (control), DHT (10nM; Sigma Aldrich), enzalutamide (10 µM, Selleckchem), or a combination of DHT and enzalutamide for 18 hours. The DNA fragments isolated by FAIRE were used for library preparation with the Roche KAPA library prep kit according to the manual and sequenced on the Illumina HiSeq 2500 to produce 50 bp single-end reads at the Genomics core (KU Leuven) and aligned using bwa 0.7.8-r455 against hg19. Duplicates were marked & Realigned using Picard 1.118. Peak calling was performed on the aligned files using MACS2 v2.1.0. MSPC v4.0.2 was used to jointly analyze the peaks called in the three replicates from each sample and to derive a common peak set across replicates. DiffBind v2.14.0 was used to explore peak overlaps and differential accessibility between samples. Read distribution analysis around transcription start sites, MYC binding sites, and AR binding sites was performed by counting the average number of reads across replicates for each sample condition in 100bp bins extending 1kb in either direction of the site. The value at the center of the resulting distributions was compared between samples using the *t*-test to assess for differences in chromatin openness at these sites.

### Software

Analyses were performed using R v3.6.3 and Python v3.7.0.

### Statistical testing

Statistical testing was performed using R v3.6.3. Statistical tests used are indicated in the text and in figure legends. The Shapiro-Wilk test was used to test for normality.

### Single-cell RNA pre-processing and quality control

The Cell Ranger output was used as the input to Seurat v3.2.0 (Butler et al., 2018; Stuart et al., 2019) for further analysis. For each sample, poor quality cells were filtered based on the number of detected genes, the total number of molecules detected, and the percentage of reads arising from the mitochondrial genome. Specific thresholds for each were adjusted per sample to preserve a maximal number of cells. To address the effects of cell cycle heterogeneity in the data, each cell was scored for its expression of genes associated with S or G2/M phases (gene sets provided within Seurat) using the Seurat CellCycleScoring function. The difference between the G2/M and S phase scores was regressed out using sctransform (Hafemeister and Satija, 2019).

### Single-cell RNA clustering

The mutual nearest neighbor approach fastMNN (Haghverdi et al., 2018) was used to integrate the four LNCaP samples using 2000 integration features and account for batch effect. Clustering and UMAP non-linear dimensionality reduction were performed using Seurat v3.2.0. The marker genes of each cluster were determined by identifying genes differentially expressed in each cluster compared to all other clusters based on the generalized linear model MAST framework v1.12.0 (Finak et al., 2015) and using the number of RNA reads as a latent variable. A gene was considered to be differentially expressed with Bonferroni corrected p-value < 0.01, at least 10% of the cells in the cluster expressing the gene, and an average log-fold change of at least 0.25.

### Cluster and sample characterization

We utilized hallmark gene sets from the Broad Institute MSigDB (Liberzon et al., 2015) to characterize clusters and samples based on their differentially expressed genes. Gene set variation analysis (GSVA) was performed using the GSVA package v1.36.2 to characterize each cluster overall using gene read counts, Poisson kernels, and the gsva method. To characterize the gene expression changes within each cluster between samples, all genes making up the integrated dataset were ranked based on their average log-fold change. The fgsea package v1.14.0 was then used to perform gene set enrichment analysis for the MSigDB hallmark gene sets using 1000 permutations. The RNA velocities of single cells in the scRNA-seq samples were assessed using scVelo v0.2.2 (Bergen et al., 2020). Differentiation states of each cell in each sample were predicted using cytoTRACE v0.3.3 (Gulati et al., 2020).

### Single-cell ATAC pre-processing and quality control

The output of the Cell Ranger ATAC pipeline was used as the input to Signac package v0.2.5 for further analysis. For each sample, poor quality cells were filtered based on the following features: strength of nucleosome binding pattern, transcription start site enrichment score as defined by ENCODE, total number of fragments in peaks, fraction of fragments in peaks, and percentage of reads in ENCODE blacklisted genomic regions. Specific thresholds for each adjusted per sample. Data normalization and dimensionality reduction was done using latent semantic indexing (LSI), consisting of term frequency-inverse document frequency (TF-IDF) normalization and singular value decomposition (SVD) for dimensionality reduction using the top 50% of peaks in terms of their variability across the samples. The first LSI component reflected sequencing depth across the samples and was not utilized in downstream analyses.

### Single-cell ATAC clustering

Integrated clustering of the scATAC-seq samples was performed with harmony v1.0 (Korsunsky et al., 2019) using LSI embeddings. The resulting harmony-adjusted cell embeddings were used as input for UMAP non-linear dimensionality reduction and clustering using default parameters and the smart local moving (SLM) algorithm for modularity optimization. A “pseudo-bulk” analysis of changes in chromatin accessibility in the scATAC-seq samples was performed by pooling the reads from all good-quality cells in each sample.

Visualization of peak overlap between samples was generated using R package ggradar v0.2. Differentially accessible regions in the clusters were identified using logistic regression with the total number of peaks as a latent variable. Differentially accessible regions were annotated with their closest gene and Reactome pathway enrichment analysis was performed on the result using R package ReactomePA v1.32.0.

### Transcription factor motif enrichment

Transcription factor motif enrichment was performed in differentially accessible regions between sample conditions utilizing R package TFBSTools v1.26.0 and JASPAR database position frequency matrices retrieved from the R JASPAR2018 data package v1.1.1. The hypergeometric test was used to test for significant enrichment of motifs, taking into account sequence characteristics of the peaks (e.g. GC-frequency). Chromatin states in scATAC-seq (as defined by the enriched TFs in differentially open chromatin regions) were connected to transcriptional outputs in the scRNA-seq by assessing for overlap between the target genes of enriched TFs and differentially expressed genes in the scRNA-seq clusters. TF target genes were obtained using the GTRD database v18.06.

### Integration of scRNA-seq datasets and scRNA- and scATAC-seq datasets using label transfer

The clusters identified from the integrated clustering of scRNA-seq from LNCaP, LNCaP-ENZ48, RES-A, and RES-B (**Figure 1I**) were queried in additional scRNA-seq samples (alternative LNCaP parental, LNCaP-ENZ168, and RES-C) (**Figure 1A**). The additional scRNA-seq samples were individually clustered and anchors were identified for each additional scRNA-seq sample and the LNCaP integrated clusters. This was done using the FindTransferAnchors function with principal component analysis (PCA). The anchors were used to transfer cluster label identifiers between the two data types using the TransferData function.

LNCaP, LNCaP-ENZ48, RES-A, and RES-B had scRNA-seq and scATAC-seq data available from each sample (**Figure 1A**). These data types were integrated using the cluster label transfer procedure as implemented in Signac v0.2.5 and Seurat v3.2.0. Each scRNA-seq sample was clustered individually and its cluster labels were projected onto the matching, individually clustered scATAC-seq sample, or vice versa. The clustering resolution of each sample was assessed and decided using clustree v0.4.3 (Zappia and Oshlack, 2018). Briefly, RNA-seq expression levels were imputed from the scATAC-seq data by defining for each gene a genomic region including the gene body and 2kb upstream of the transcription start site and taking the sum of scATAC-seq fragments within the region. Anchors were identified for condition-matched scRNA- and scATAC-seq samples using the FindTransferAnchors function and canonical correlation analysis (CCA) was performed on the scRNA expression values and the scATAC imputed gene expression values. The anchors were used to transfer cluster label identifiers between the two data types using the TransferData function.

### Gene sets and clinical data analysis

Each gene signature or set was assessed for enrichment and scored in a sample using the GSVA package v1.36.2. In cases where the expression of a gene set was assessed at the single-cell level, the GetModuleScore function in Seurat was used to generate an average expression score per cell. Survival analyses were performed using the survival package v3.2-3 and Kaplan-Meier curves were plotted using the survminer package v0.4.8. For single signature survival analyses, median GSVA score was used to stratify patients into low and high expressing groups for the signature. For survival analyses of multiple signatures, samples were clustered using their GSVA enrichment scores for each signature using Euclidean distance and hierarchical clustering. The clustering result was then used to define the two-group split of samples for the survival analysis. To generate the PROSGenesis signature, we extracted the gene expression profile associated with the regenerative mouse prostate luminal 2 cells reported in Karthaus *et al* and found 78 genes with homologues in humans that have been profiled in our scRNA-seq dataset.

### Spatial transcriptomics analysis of primary prostate cancer tissue

Two sections of cryopreserved prostate cancer tissue from one patient (pT=2b, T1c, Gleason 6, PSA 3.5 ng/mL) were profiled for spatial transcriptomics using the Visium Spatial library preparation protocol from 10x Genomics with a resolution of 55 µm (1-10 cells) per spot. The tissues were cryosectioned at 10 µm thickness to Visium library preparation slide, fixed in ice-cold 100% methanol for 30 min, H&E stained with KEDEE KD-RS3 automatic slide stainer and the whole-slide was imaged using Hamamatsu NanoZoomer S60 digital slide scanner.

Sequencing library preparation was performed according to Visium Spatial Gene Expression user guide (CG000239 Rev D, 10x Genomics), using 24 min tissue permeabilization time. Sequencing was done on the Illumina NovaSeq PE150 sequencer at Novogene Company Limited, Cambridge, UK’s sequencing core facility, aiming at 50,000 read pairs per tissue covered spot.

Sequenced data was first processed using Space Ranger v1.2 from 10x Genomics to obtain per-spot expression matrices for both sections. Downstream processing and clustering was then performed using Seurat v3.2.0. Normalization of the data was performed with sctransform to account for differences in sequencing depth across spots. Clustering was performed using the FindClusters function using a resolution parameter value of 0.8. The resulting clusters were found to correspond to histological characteristics of the tissue. The GetModuleScore function of Seurat was used to score the spots for our scRNA-seq derived gene signatures, as well as length-matched random housekeeping gene signatures from the Housekeeping and Reference Transcript Atlas (Hounkpe et al., 2021). The distributions of the gene expression scores for the housekeeping gene sets and our scRNA-seq signatures were compared to determine the 90th percentile as a score cutoff at which we considered a spot to have high expression of the scRNA-seq signature, allowing for 5% false positives (spots scoring above the threshold for housekeeping gene sets).

## Supporting information

See also

## Acknowledgments

We thank the Genomics Core Facility at Institute for Cancer Research, OUH, for the support during preparation of the scATAC- and RNA-seq and the Tampere University histology core facility and Sari Toivola for skillful assistance during Visium experiments. The study was financially supported by the Finnish Cultural Foundation (ST), Academy of Finland (#312043 (MN), #310829 (MN), #324009 (KK), #328928 (KK), #333545 (KG)), Cancer Foundation Finland (MN, KK, KG), Sigrid Jusélius Foundation (MN, KK, KG), Emil Aaltonen Foundation (KG), Finnish Cancer Institute (MN), Norwegian Cancer Society (#198016-2018)(AU, NE), The Norman Jaffe Professorship in Pediatrics Endowment Fund (SC), MD Anderson Colorectal Cancer Moon Shot Program (SC), Oncode Institute (SP), Finnish Cultural Foundation North Savo Regional fund (KK, RK), University of Eastern Finland Doctoral Programme in Molecular Medicine (RK), K. Albin Johansson Foundation (KK), John Black Foundation (IM), Human Cell Atlas Seed Network - Retina (WW), Chan Zuckerberg Institute (WW), NIH R01CA183793 (WW, SC), NIH R01CA239342 (WW), NIH R01CA158113 (WW), P30CA016672 (WW), the Fonds Wetenschappelijk Onderzoek-Vlaanderen (GOA9816N)(FC), KU Leuven (C14/19/100)(FC), Kom op tegen Kanker (KOTK)(FC). WD is holder of a fellowship of KOTK. The results published here are in part based upon data generated by The Cancer Genome Atlas project established by the NCI and NHGRI.

## Notes

### Competing Interest Statement

The authors have declared no competing interest.

### Summary of Updates

We added the authors Tolonen, Hakkinen, Erickson, and Lamb to the list of authors

