## Supplementary material for "Single-cell ATAC and RNA sequencing reveal pre-existing and persistent subpopulations of cells associated with relapse of prostate cancer": See also

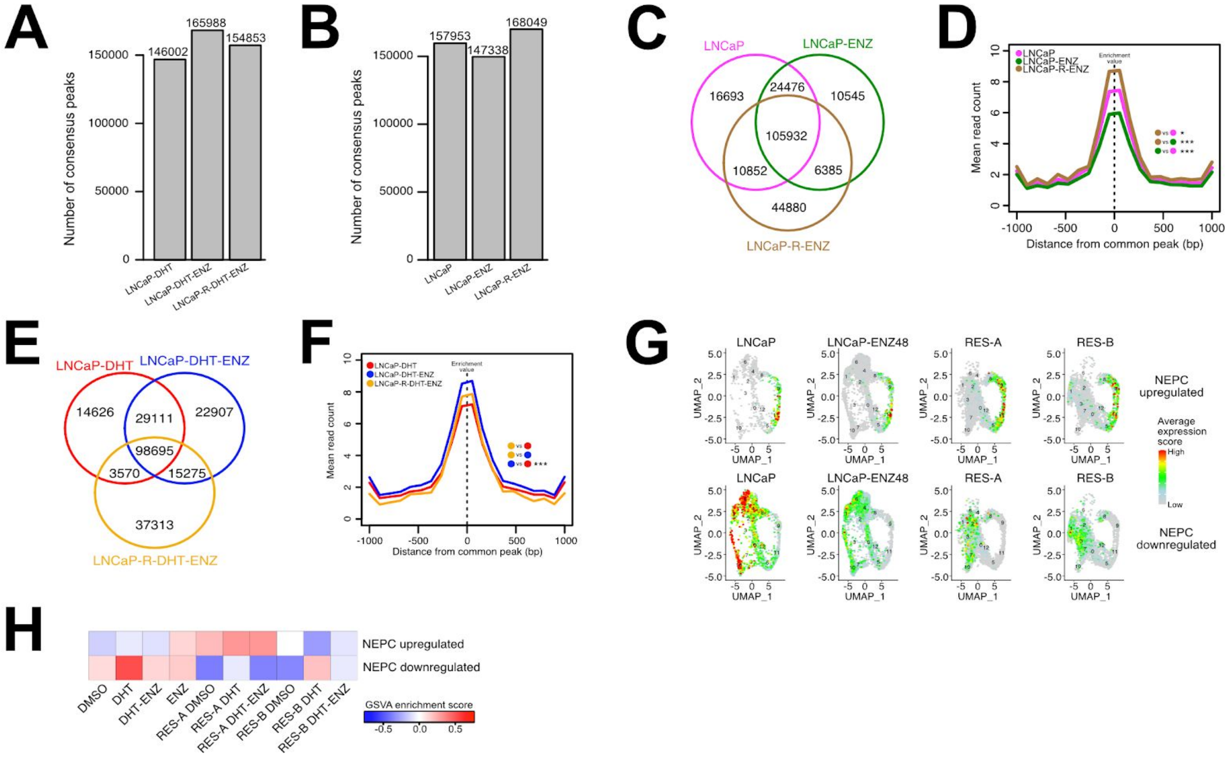


**Figure S1 (Relative to Figure 1). (A-B)** Barplots of consensus FAIRE-seq peaks in LNCaP samples treated with androgens or in castrate conditions. **(C)** Venn diagram of FAIRE-seq peaks shared or unique to LNCaP samples in androgen-deprived conditions. **(D)** Mean read count distribution within a 2kb interval around FAIRE-seq sites shared by all LNCaP samples in androgen-deprived conditions. **(E)** Venn diagram of FAIRE-seq peaks shared or unique to LNCaP samples in castrate conditions. **(F)** Mean read counts observed within a 2000bp window of peaks shared by all LNCaP samples in castrate conditions. **(G)** Feature plot showing the average expression score of each cell in the four LNCaP scRNA-seq samples for the NEPC up- and downregulated gene sets. Red indicates high expression and grey indicates low expression. **(H)** GSVA enrichment scores for NEPC up- and downregulated genes in bulk RNA sequencing of LNCaP exposed to various treatments.

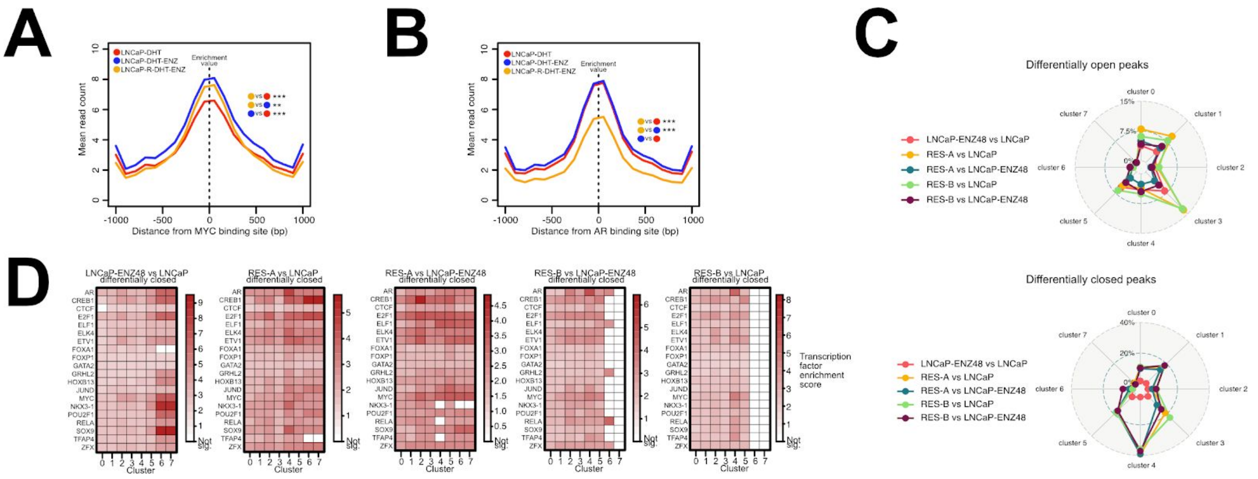


**Figure S2 (Relative to Figure 2). (A-B)** Mean read counts for LNCaP samples in androgen-exposed conditions within a 2kb window around **(A)** MYC binding sites and **(B)** around AR binding sites. **(C)** Radar plots of open and closed differentially accessible regions (DARs) in pairwise sample comparisons, expressed in percentage of the total possible regions in the respective scATAC-seq cluster. **(D)** Prostate cancer-associated transcription factor motif enrichments in closed DARs in pairwise sample comparisons. Enrichments with a Benjamini-Hochberg method adjusted hypergeometric test p-value < 0.05 are shown in colors, while non-significant enrichment are shown in white.

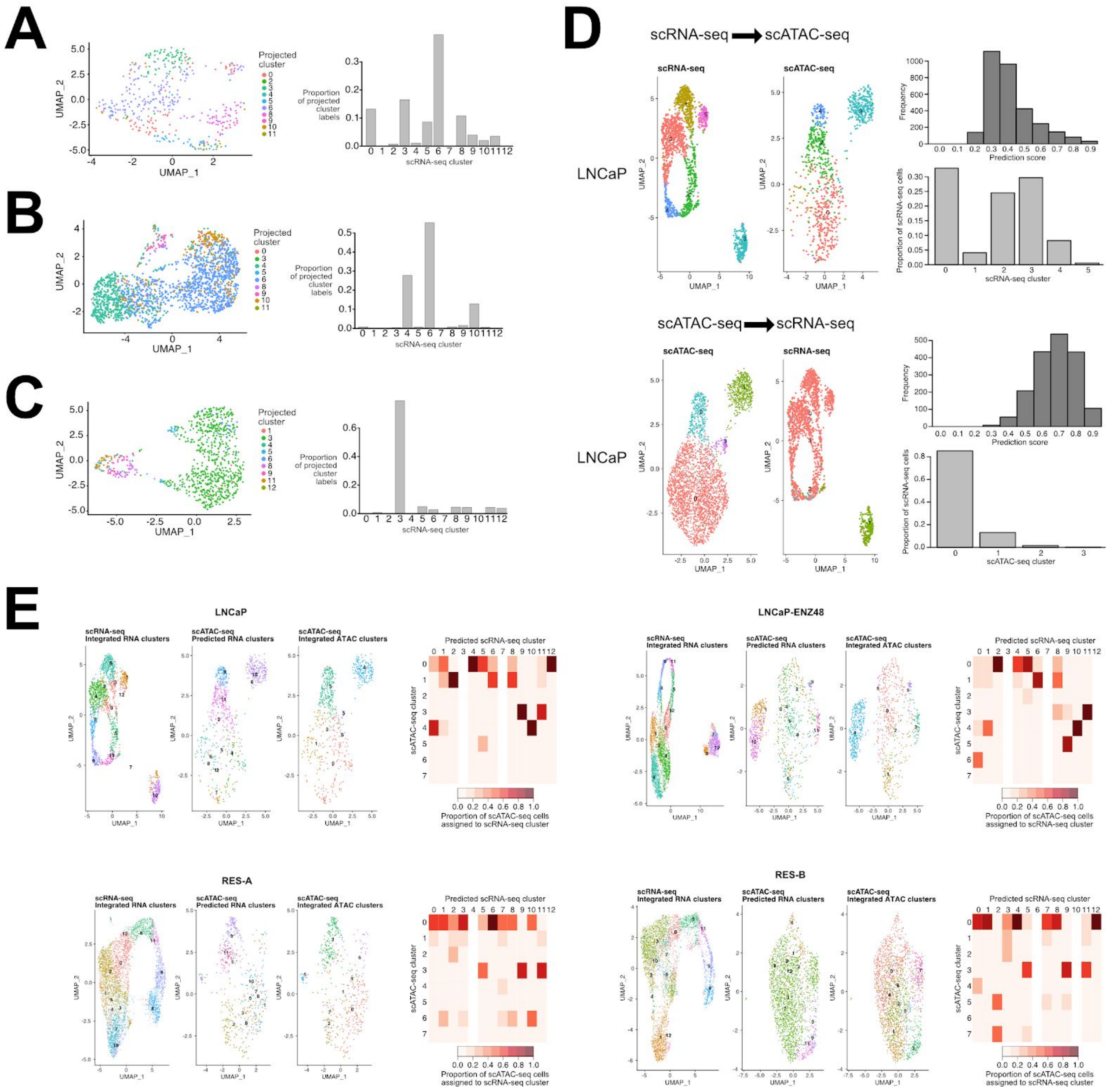


**Figure S3 (Relative to Figure 3). (A)** Cluster label transfer from integrated clustering of the scRNA-seq data to an enzalutamide resistant LNCaP sample treated with ENZ for 9 months (RES-C). Each cell is colored according to the scRNA-seq cluster that it is predicted to belong to. The barplot shows the proportion of the projected cluster labels for each scRNA-seq cluster. **(B)** Cluster label transfer from integrated clustering of the scRNA-seq data to an LNCaP sample treated with DMSO from an outside dataset. Each cell is colored according to the scRNA-seq cluster that it is predicted to belong to. The barplot shows the proportion of the projected cluster labels for each scRNA-seq cluster. **(C)** Cluster label transfer from integrated clustering of the scRNA-seq data to a scRNA-seq LNCaP sample treated with ENZ for 168 hours (LNCaP-ENZ168). For the LNCaP-ENZ168 sample, each cell is colored according to the scRNA-seq cluster that it is predicted to belong to. The barplot shows the proportion of the projected cluster labels for each scRNA-seq cluster. **(D)** Sample-wise transfer of cluster labels from scRNA-seq to scATAC-seq, and from scATAC-seq to scRNA-seq, exemplified in parental LNCaP. For each cluster label transfer direction, the confidence scores of the assigned cluster labels for each cell in the query are plotted as a histogram, and the proportion of query cells assigned to each cluster from the reference is shown as a barplot. In the top figure set, the scRNA-seq clusters are used as the reference and projected onto the scATAC-seq cells. In the scATAC-seq UMAP, cells that could be assigned a cluster label from the scRNA-seq with a confidence score of 0.5 or higher are colored according to their predicted scRNA-seq cluster assignment. In the bottom figure set, the cluster label transfer process is performed by using the scATAC-seq clusters as the reference and projecting the cluster labels onto the scRNA-seq cells. **(E)** Sample-wise transfer of cluster labels from scRNA-seq to scATAC-seq for the integrated scRNA-seq clusters as shown in **Figure 3A**. For each sample condition, the scRNA-seq sample was clustered individually and the cells labeled according to their integrated clusters. These clusters were then queried in the scATAC-seq cells and those with label transfer confidence scores > 0.4 were labeled according to their predicted scRNA-seq cluster. The same scATAC-seq cells were finally labeled according to their integrated scATAC-seq cluster to visualize the matching cell states between the data types. For each sample condition, a matrix is shown to depict the proportion of scATAC-seq cells for the sample condition assigned to each scRNA-seq cluster. The proportions were calculated for each scRNA-seq cluster, with the total as the number of cells from the scATAC-seq that could be confidently assigned to an scRNA-seq cluster (confidence score > 0.4).

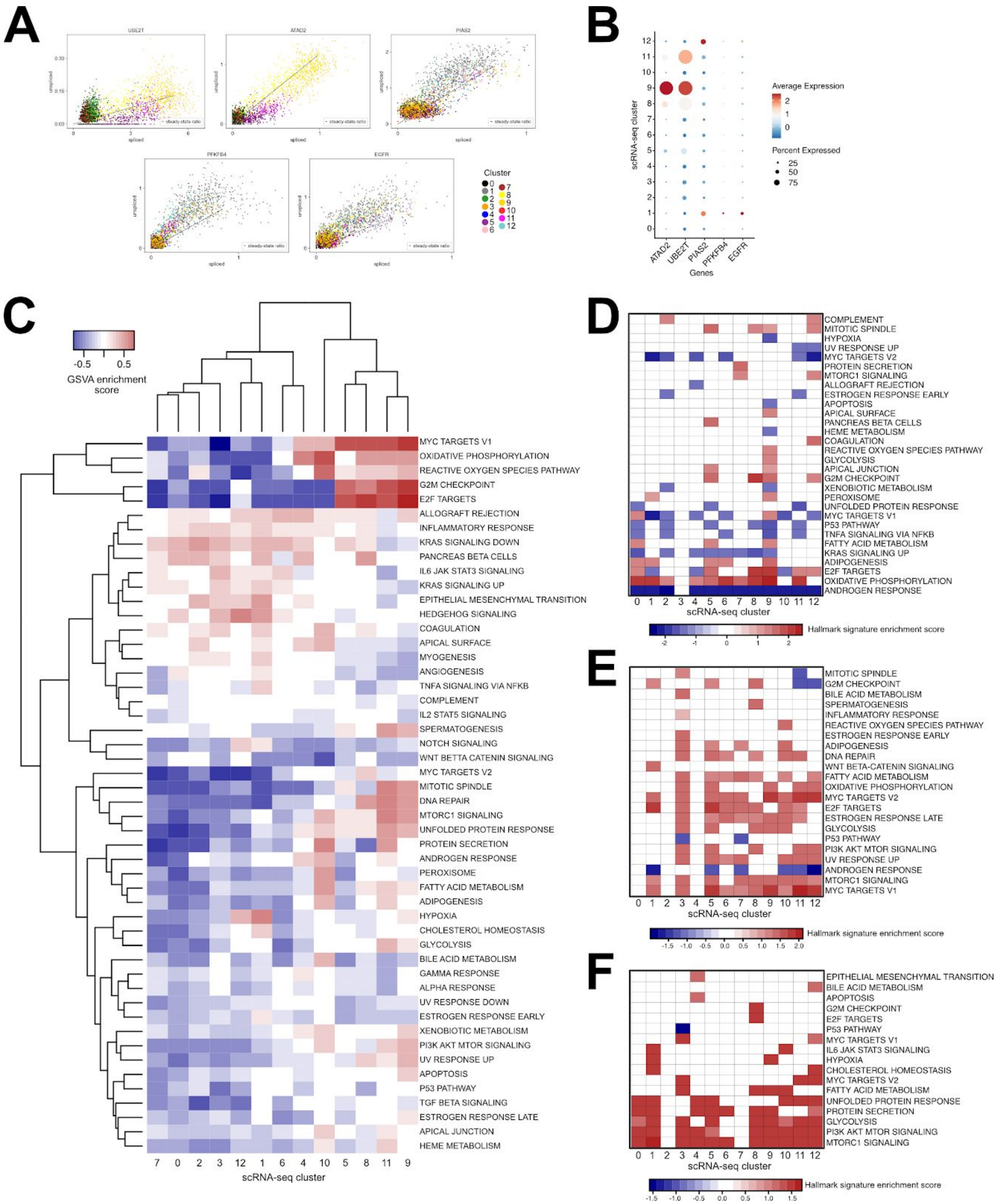


**Figure S4 (Relative to Figure 4). (A)** Examples of marker genes for RNA velocity across clusters identified from scRNA-seq for RES-A and RES-B. For each gene the abundance of spliced versus unspliced mRNA is shown, with the black dashed line indicating equal amounts of spliced and unspliced mRNA. Clusters of cells identified from scRNA-seq are shown in different colors. **(B)** Expression of RNA velocity marker genes in the clusters identified from scRNA-seq shown as a dot plot. **(C-F)** Gene set enrichment analysis of MSigDB hallmark gene sets in scRNA-seq. Heatmap colors correspond to normalized GSEA enrichment scores with Benjamini-Hochberg adjusted p-values $< 0.05$. Hallmark gene sets with adjusted p-values $> 0.05$ are shown in white. **(C)** Hallmark gene set enrichments for the average gene expression profile of each scRNA-seq cluster presented as a clustered heatmap. Hallmark gene set enrichments for changes in gene expression between **(D)** LNCaP-ENZ48 and LNCaP, **(E)** RES-A and LNCaP, and **(F)** RES-B and LNCaP.

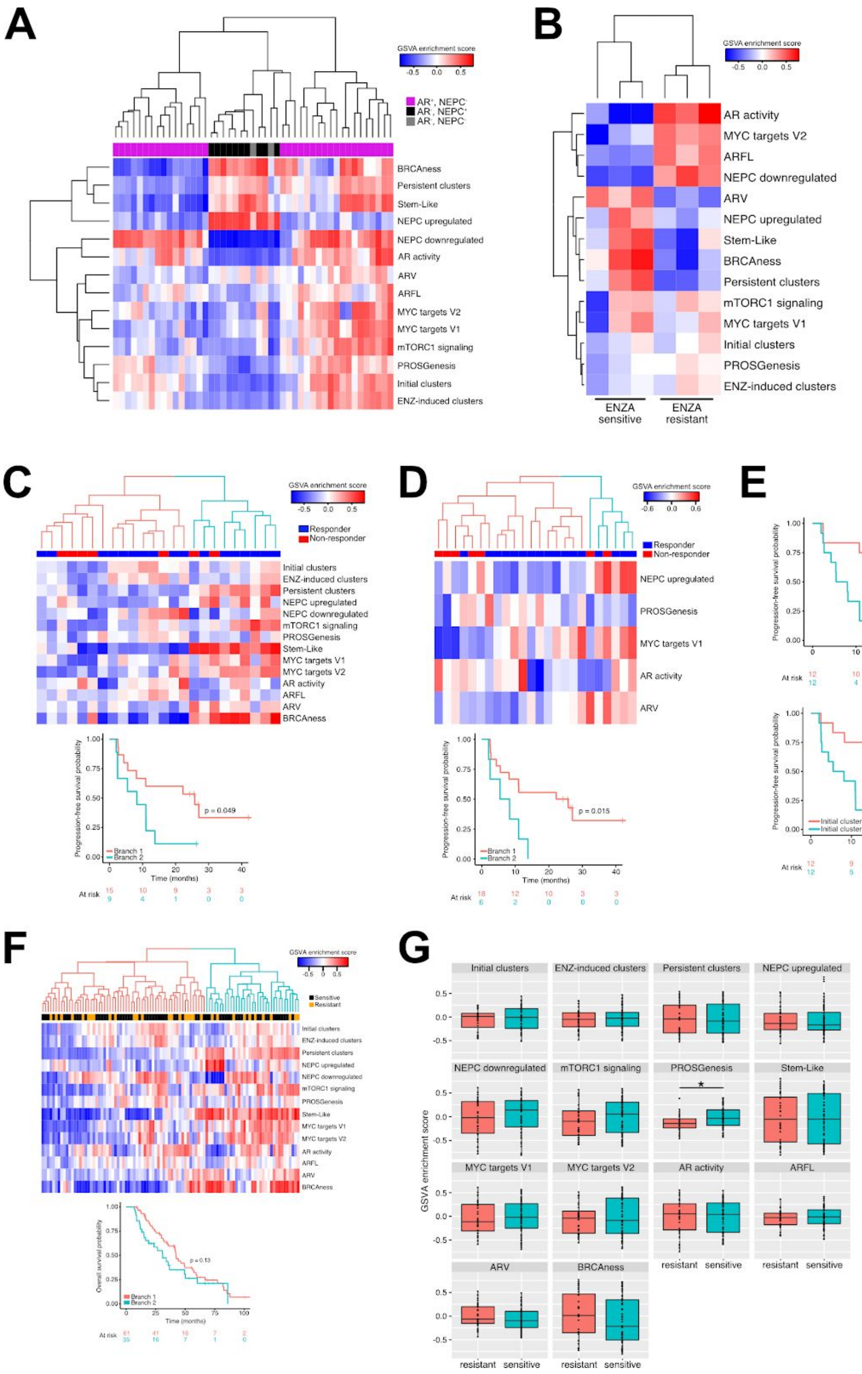


**Figure S5 (Relative to Figure 5). (A)** Heatmap of GSVA enrichment scores for a dataset consisting of tumors of varying AR and NEPC status. **(B)** Heatmap of GSVA enrichment scores for ENZ-sensitive and ENZ-resistant xenografts. **(C)** GSVA enrichment scores for all single-cell gene signatures in Alumkal *et al*, with hierarchical clustering of the patients into two groups based on the scores indicated in pink and cyan in the column dendrogram. Kaplan-Meier curve with patients stratified into two groups determined by the clustering. Log-rank p-value is indicated above the curve. **(D)** GSVA enrichment scores for patients from Alumkal *et al* for the five gene signatures selected via stepwise variable selection to form the best Cox proportional hazards model. Kaplan-Meier curve for two groups of patients identified via hierarchical clustering of the GSVA scores. Log-rank p-value is indicated above the curve. **(E)** Kaplan-Meier curves for Alumkal *et al* patients stratified into two groups based on median GSVA score for the PROSGenesis and initial cluster signatures. Log-rank p-value is indicated above the curves. **(F)** Single-cell-derived gene sets cannot be used to identify patients with shorter overall survival in the SU2C West Coast Dream Team dataset from Quigley *et al.* Heatmap of GSVA enrichment scores for patients from Quigley *et al.* Hierarchical clustering of the patients into two groups is shown by the pink and cyan colors in the column dendrogram. This patient grouping was used to stratify patients for the overall survival Kaplan-Meier curve. Log-rank p-value is indicated above the curve. **(G)** Boxplots of GSVA enrichment scores for ENZ-resistant and sensitive patients from the SU2C West Coast Dream Team dataset from Quigley *et al*. Pink boxplots indicate the GSVA scores of ENZ-resistant samples and cyan boxplots show the GSVA scores of ENZ sensitive samples. Differences in GSVA scores for signatures between ENZ sensitive and resistant samples were assessed using Wilcoxon rank-sum test, with the PROSGenesis showing a significant difference ($p = 0.024$, indicated with asterisks).

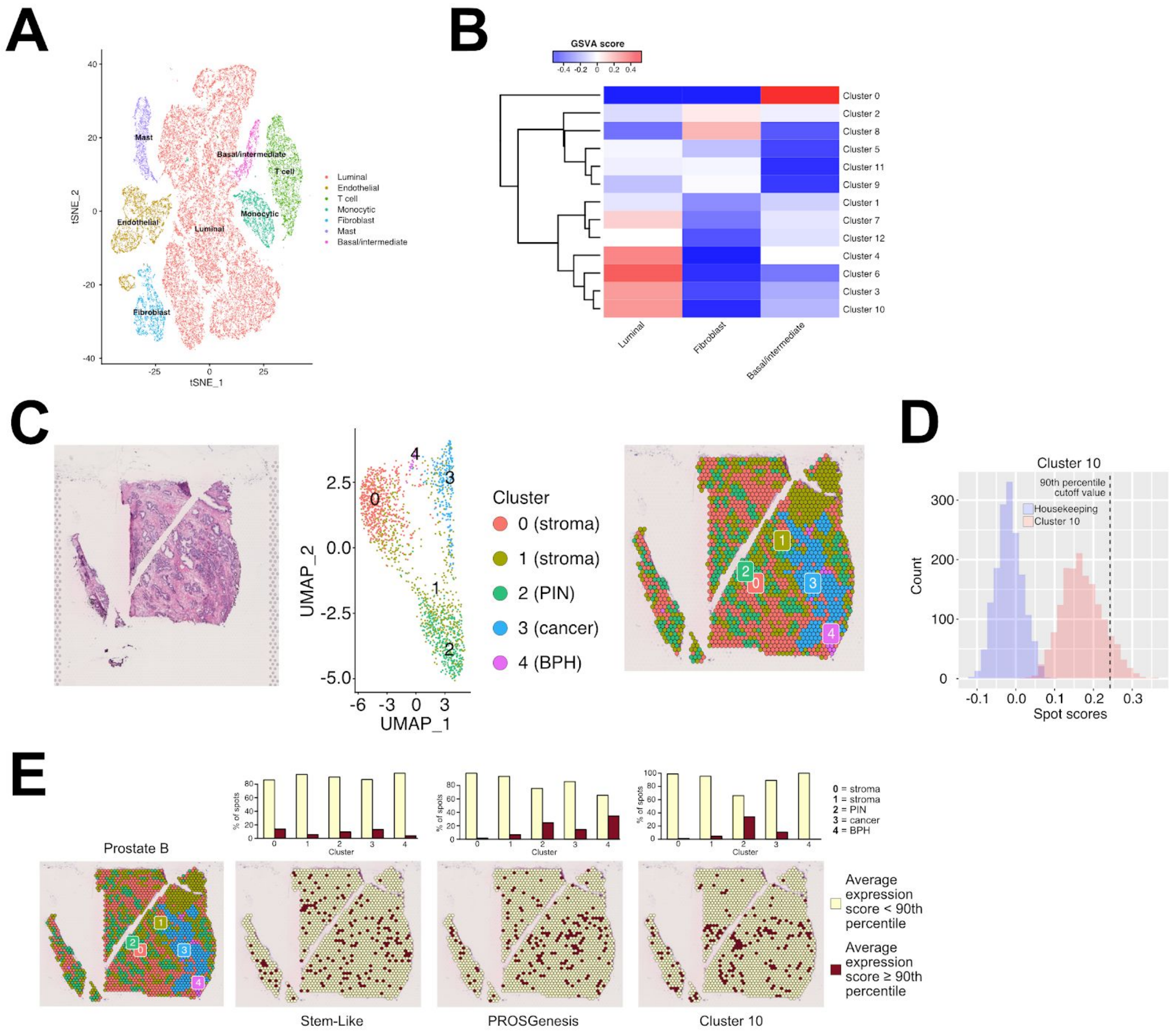


**Figure S6 (Relative to Figure 6). (A)** tSNE visualization of the cell types of the cells from Chen *et al* (Chen et al., 2021). **(B)** GSVA enrichment scores for LNCaP scRNA-seq individual cluster signatures in luminal, basal/intermediate, and fibroblast cells from Chen *et al*. GSVA enrichment scores were generated from the average expression profile of each cell type. **(C-E)** Spatial transcriptomics from a prostate cancer tissue section, Prostate B. **(C)** The leftmost panel shows the H&E staining of the tissue section. In the middle, the UMAP visualization of the clusters of the spots on the spatial transcriptomics slide. Each cluster is also labeled according to its histological tissue type, with clusters 0 and 1 corresponding to stroma, cluster 2 corresponding to prostatic intraepithelial neoplasia (PIN), cluster 3 corresponding to the prostate adenocarcinoma, and 4 corresponding to benign prostatic hyperplasia (BPH). The rightmost panel shows the UMAP clusters of spots overlaid on the H&E slide. **(D)** Sensitivity analysis of Cluster 10 gene signatures scores in ST against the score distribution of a control housekeeping gene signature (see **Methods**). **(E)** The leftmost panel shows the UMAP clusters of spots overlaid on the H&E slide. Each spot was scored according to its expression of genes in the

Stem-Like, PROSGenesis, and cluster 10 signatures. For each signature, spots scoring at or above the 90th percentile ("high") are colored in red, while spots scoring below the 90th percentile ("low") are colored in yellow. The barplots indicate the percentage of spots in each cluster scoring high or low for each signature.

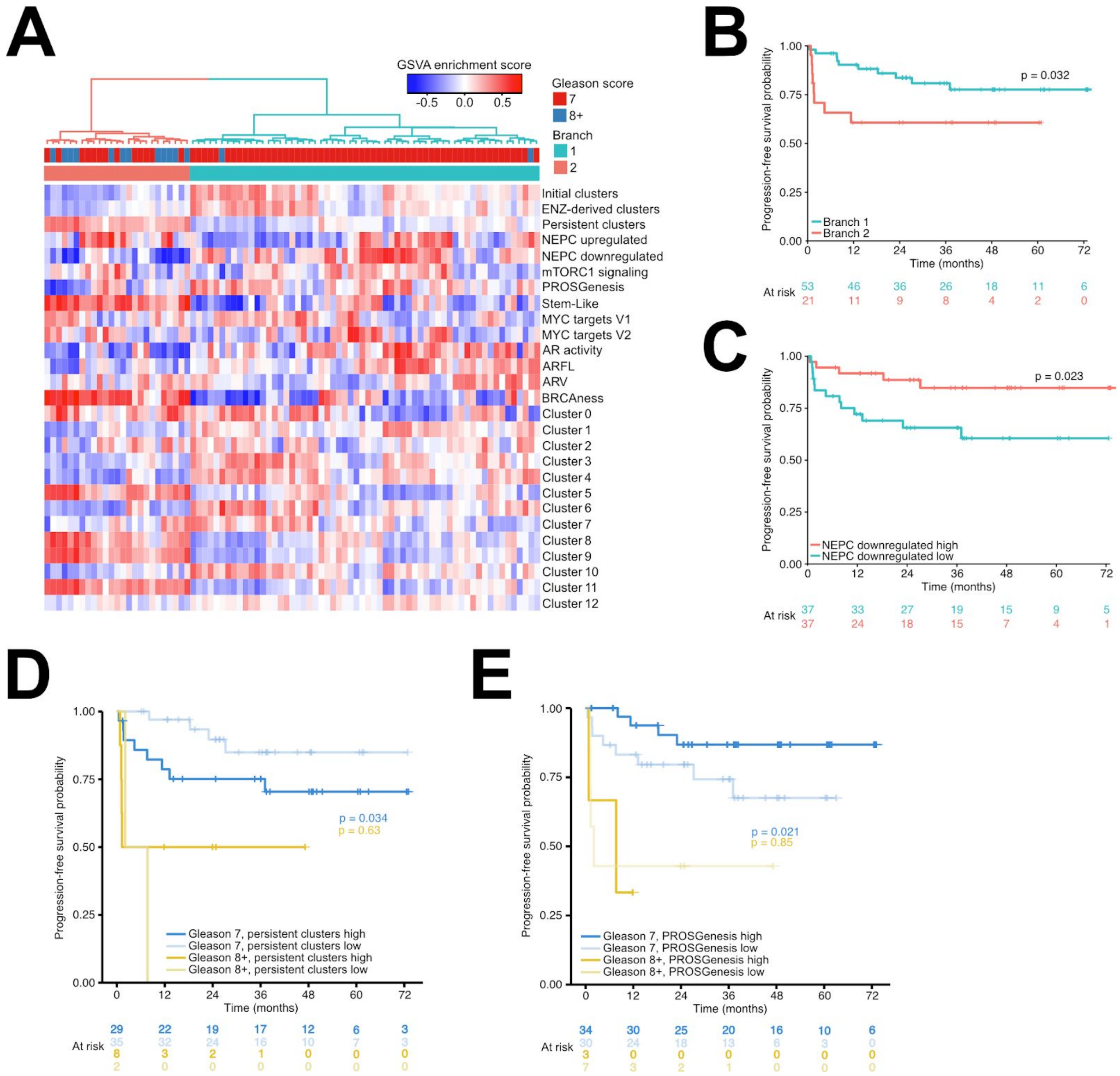


**Figure S7 (Relative to Figure 7). (A)** Heatmap of GSVA enrichment scores for all single-cell-derived gene signatures in the ICGC-EOPC cohort, including the marker gene set for each scRNA-seq cluster. Hierarchical clustering of the GSVA scores was used to separate the samples into two groups, marked Branch 1 and Branch 2. **(B)** Kaplan–Meier survival curve for ICGC-EOPC patients stratified into two groups as indicated in Panel B. **(C)** Kaplan-Meier survival curves for ICGC-EOPC patients stratified into two groups based on median GSVA score for NEPC downregulated gene signature. **(D-E)** Kaplan-Meier curves for ICGC-EOPC patients stratified into four groups based on Gleason score and median GSVA score for the persistent cluster signature or the PROSGenesis signature. Log-rank p-values are shown within the plots.
